# Twinfilin uncaps filament barbed ends to promote turnover of lamellipodial actin networks

**DOI:** 10.1101/864769

**Authors:** Markku Hakala, Hugo Wioland, Mari Tolonen, Antoine Jegou, Guillaume Romet-Lemonne, Pekka Lappalainen

## Abstract

Coordinated polymerization of actin filaments provides force for cell migration, morphogenesis, and endocytosis. Capping Protein (CP) is central regulator of actin dynamics in all eukaryotes. It binds actin filament (F-actin) barbed ends with high affinity and slow dissociation kinetics to prevent filament polymerization and depolymerization. In cells, however, CP displays remarkably rapid dynamics within F-actin networks, but the underlying mechanism has remained enigmatic. We report that a conserved cytoskeletal regulator, twinfilin, is responsible for CP’s rapid dynamics and specific localization in cells. Depletion of twinfilin led to stable association of CP with cellular F-actin arrays and its treadmilling throughout leading-edge lamellipodium. These were accompanied by diminished F-actin disassembly rates. In vitro single filament imaging approaches revealed that twinfilin directly promotes dissociation of CP from filament barbed ends, while allowing subsequent filament depolymerization. These results uncover an evolutionary conserved bipartite mechanism that controls how actin cytoskeleton-mediated forces are generated in cells.

---

Coordinated polymerization of actin filaments (F-actin) generates pushing force, which drives the protrusion of lamellipodia at the leading edge of migrating cells, and contributes to the formation of plasma membrane invaginations in endocytic processes^1–4^. A large array of actin-binding proteins regulates the dynamics and organization of actin filaments in these processes, but only few (<10) of them are conserved in evolution from protozoan parasites to animals^5^. Among these ‘core’ actin-binding proteins are the heterodimeric Capping Protein (CP) and twinfilin.

Generation of membrane protrusions requires that a subset of actin filament barbed ends is capped to funnel the assembly-competent actin monomers to a limited number of growing barbed ends^6–8^. CP is the most prominent actin filament barbed end capping protein in most organisms and cell types^9, 10^. It is an essential component of *in vitro* reconstituted actin-based motility system^11^ and actin-based processes in cells^12–14^. Moreover, its inhibition in animal non-muscle cells disturbs actin-based lamellipodial morphology, protrusion and cell migration^15–18^. In addition, CP plays an important role in controlling the length and the density of branches within actin filament networks nucleated by the Arp2/3 complex^19^.

The activities of CP are controlled by several proteins^9, 10^. V-1/ myotrophin binds and sequesters CP with nanomolar affinity and inhibits its capping activity^20–23^. Moreover, the capping protein interaction (CPI)-motif containing proteins, such as CARMILs (capping protein, Arp2/3 and myosin-I linker protein)^24^, interact with CP to reduce its affinity to both actin filament barbed ends^25–27^ and to V-1^22^ through an allosteric mechanism^25, 28^. Depletion of CARMILs and other CPI motif containing proteins disrupts the subcellular localization of CP^29–33^. Thus, it was suggested that CPI-motif proteins activate CP near the plasma membrane by competing with V-1^22^.

CP binds actin filament barbed ends with sub-nanomolar affinity and displays very slow dissociation kinetics (half-time ~30 minutes)^34–36^. However, its turnover in cells is orders of magnitude faster^37, 38^ and the localization is restricted to very distal edge in lamellipodia of motile cells^16, 37^. The discrepancy of data between biochemical and cellular experiments suggests that additional factors might regulate uncapping of CP-capped barbed ends. Both CARMIL and ADF/cofilin can enhance uncapping of filament barbed ends in vitro^26,36,39^. However, to our knowledge the evidence of filament uncapping in cells is missing.

Twinfilin is a conserved ADF/cofilin-like protein that is composed of two actin-binding ADF-H (actin depolymerization factor homology) domains, followed by CPI-motif containing C-terminal tail^40^, which binds CP with high affinity^41–43^ and interacts with membrane phosphoinositides^42^. Earlier biochemical studies demonstrated that twinfilin can sequester actin monomers^44–46^, cap filament barbed ends^47^ and accelerate actin filament depolymerization *in vitro* in concert with cyclase-associated protein (Srv/CAP)^48, 49^. Importantly, *in vivo* studies show that twinfilin contributes to various actin-dependent processes^44,50–53^. For example, depletion of twinfilin in Drosophila S2 cells lead to expansion of the lamellipodium, and budding yeast cells lacking twinfilin display defects in the turnover of endocytic actin patches^17, 44^. However, whether twinfilin contributes to actin-dependent processes by functioning as an actin monomer sequestering, filament capping, or filament depolymerizing protein, or through yet another unknown mechanism, has remained enigmatic. The presence of two twinfilin genes in mammals, that encode the ubiquitously expressed twinfilin-1 and twinfilin-2a, and a muscle specific isoform twinfilin-2b^40, 54^, further complicates the analysis of twinfilin’s functions in cells.

Here we reveal that twinfilin controls actin filament disassembly in cells by accelerating the dissociation of CP from filament barbed ends. By utilizing twinfilin-1/twinfilin-2 double knockout cell lines, we show that the correct localization and dynamics of CP in cells are dependent on twinfilin. Moreover, our measurements of single actin filament dynamics in vitro revealed the molecular mechanism by which CP rapidly dissociates from the actin filament barbed in presence of twinfilin. Together, our data explain how CP is enriched at the distal edge of lamellipodial actin filament networks and undergoes rapid turnover at actin filament barbed ends in cells.

## Results

### Depletion of twinfilin results in abnormal actin filament accumulation and lamellipodial dynamics

To reveal the role of twinfilin in actin dynamics in mammalian cells, we generated *twinfilin-1* and *twinfilin-2* knockout mouse melanoma B16-F1 cell lines as well as *twinfilin-1/twinfilin-2* double knockout cell lines (hereafter referred to as twf1/twf2-KO) by utilizing CRISPR/Cas9 system (Supplementary figure 1A, C-F). Inactivation of either *twinfilin-1* or *twinfilin-2* gene did not result in drastic defects in the organization of the actin cytoskeleton (Supplementary figure 2A-C), suggesting that twinfilin-1 and twinfilin-2a/b may be functionally redundant in cells. Accordingly, twf1/twf2-KO cells exhibited more severe abnormalities in actin-dependent processes. In comparison to wild-type B16-F1 cells, lamellipodia of twf1/twf2-KO cells were less smooth (Figure 1A, Supplementary figure 2A), and the velocities of lamellipodial protrusions were slower (Figure 1C, Supplementary figure 3B-C, and Supplementary movie 1). Moreover, twf1/twf2-KO cells displayed elevated accumulation of F-actin to Arp2/3-positive patches, which were enriched at the perinuclear region (Supplementary figure 2C, Supplementary figure 3A). These phenotypes were reflected by increased overall F-actin levels in twf1/twf2-KO cells, while the total levels of actin appeared unaltered compared to the wild-type cells (Figure 1B, Supplementary figure 1B, Supplementary figure 2B). Importantly, these phenotypes could be rescued by transient expression of EGFP-twinfilin-1 in twf1/twf2-KO cells, suggesting that they do not result from off-target effects (Figure 1B-C, Supplementary figure 4).

**Figure 1.**
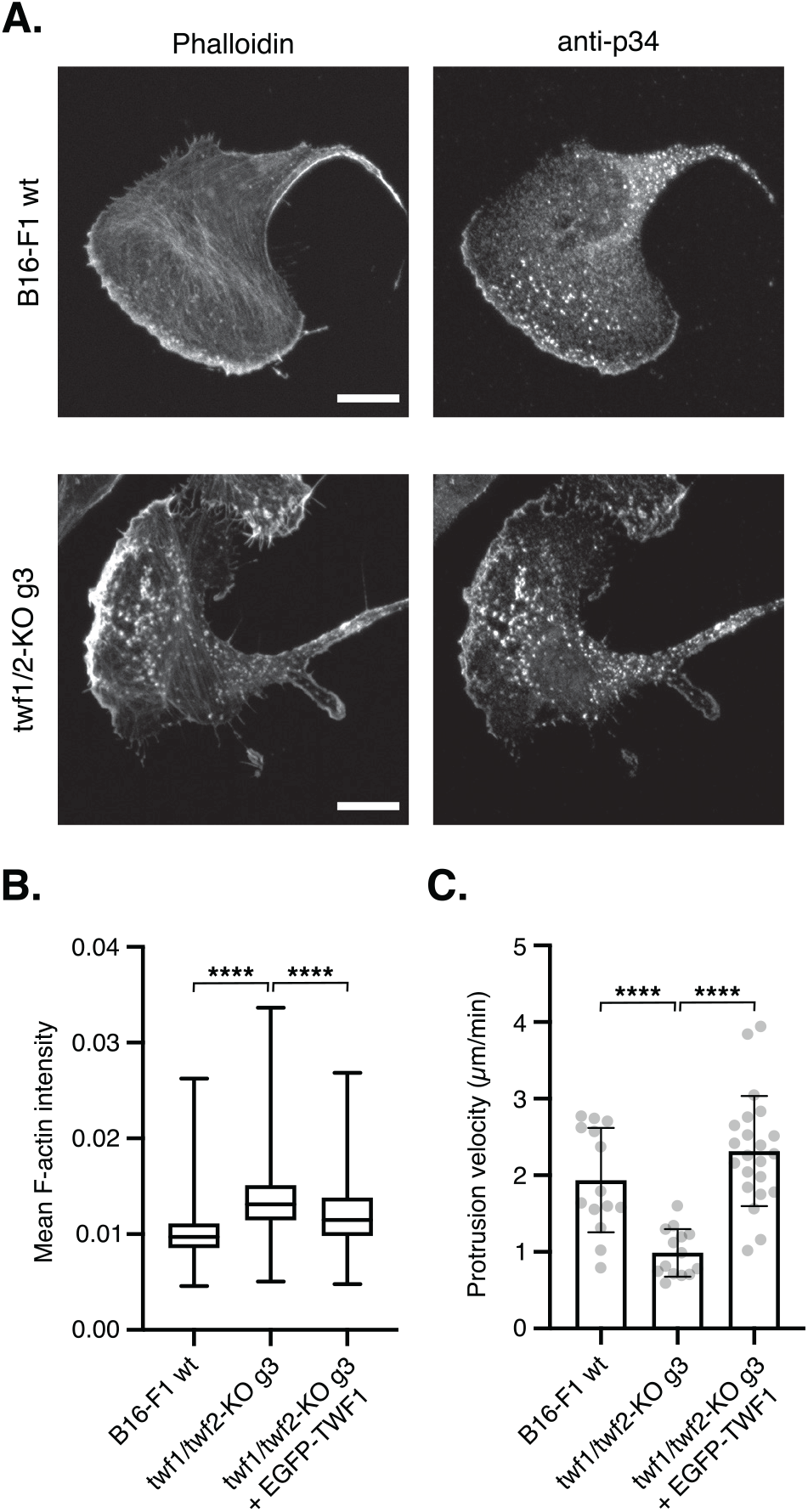
Knockout of twinfilins leads to abnormal F-actin accumulation in lamellipodia and perinuclear region. (A) Representative images of wild-type and twf1/twf2-KO mouse B16-F1 cells stained with AlexaFluor-568 phalloidin and anti-p34 antibody to visualize F-actin and the Arp2/3 complex, respectively. Scale bar = 10 *μ*m. (B) F-actin intensities in the cytoplasmic regions of wild-type, twf1/twf2-KO, and knockout cells expressing EGFP-TWF-1 measured by high-content image analysis. Number of cells analyzed were: B16-F1 wt = 4,875, twf1/twf2-KO-g3 = 5,731, twf1/twf2-KO-g3 + EGFP-TWF-1 = 197. (C) Lamellipodia protrusion velocities of wild-type, twf1/twf-2 knockout, and knockout cells expressing EGFP-TWF-1. Data represent individual cells with mean and standard deviations shown. Statistical significances in panels B and D were calculated with Mann-Whitney two-tailed test. ****, p<0.0001.

Earlier work showed that twinfilins co-localize with actin in transferrin-positive endosomes^47^. We thus examined if the F-actin-rich patches in twf1/twf2-KO cells are endocytic structures. By labelling endosomes with fluorescent transferrin, we observed that F-actin accumulated to transferrin-positive punctae especially at the perinuclear region of twf1/twf2-KO cells, whereas wild-type B16-F1 cells displayed more uniform F-actin staining (Supplementary figure 5A-B). Moreover, high-content analysis revealed a significant increase in the F-actin intensity on transferrin positive endosomes of twf1/twf2-KO cells compared to wild-type cells (Supplementary figure 5C). Together, these results provide evidence that twinfilin-1 and twinfilin-2 display redundant roles in controlling actin dynamics in lamellipodia and endosomal structures.

### Twinfilin promotes actin filament disassembly at lamellipodium

We hypothesized that the abnormal accumulation of actin filaments in twf1/twf2-KO cells is either due to increased F-actin assembly or decreased disassembly. To address this question, we applied fluorescence recovery after photobleaching (FRAP) assay to study F-actin treadmilling, and fluorescence decay after photoactivation assay to study F-actin disassembly in the lamellipodia of wild-type and twf1/twf2-KO cells. FRAP assay on B16-F1 cells expressing EGFP-β-actin revealed that the rate of F-actin treadmilling was diminished in twf1/twf2-KO cells compared to wild-type B16-F1 cells (Figure 2A-D, Supplementary movie 2). The slower F-actin treadmilling rate at the lamellipodia is consistent with the decreased lamellipodial protrusion velocity in twinfilin-deficient cells (Figure 1C).

**Figure 2.**
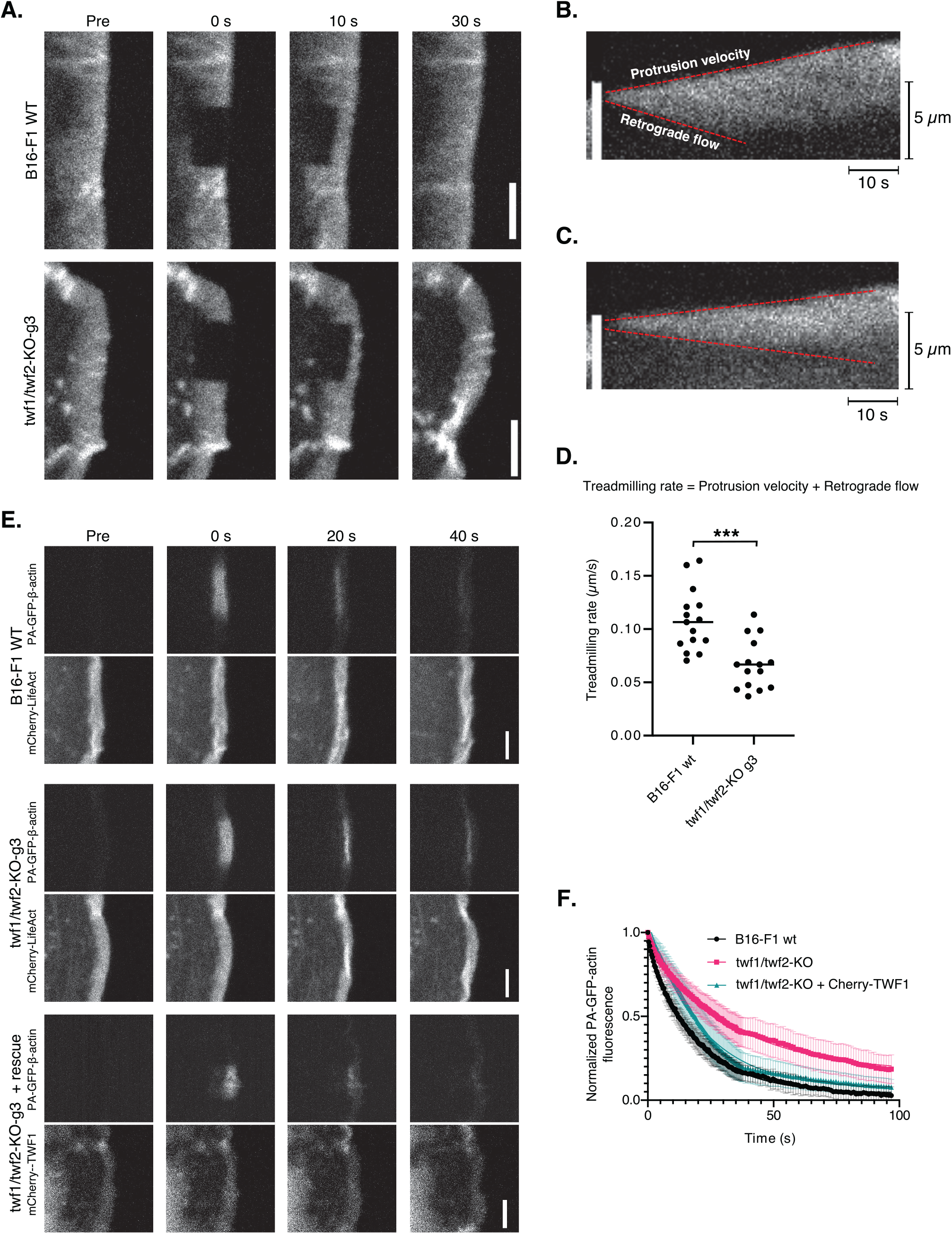
Twinfilin regulates actin dynamics in lamellipodia. (A) Fluorescence recovery after photobleaching (FRAP) of EGFP-actin in wild-type and twf1/twf2-KO B16-F1 cells. Timepoint “Pre” is a frame before bleaching, and 0 s the first frame after bleaching. Scale bar = 5 *μ*m. (B-C) examples of kymographs created from the center of the bleached region in lamellipodia. Treadmilling rates were measured as a sum of protrusion velocity and retrograde flow. (D) Treadmilling rates of EGFP-β-actin in the lamellipodia of wild-type and twf1/twf2-KO B16-F1 cells. Data represent individual experiments, and the mean values are indicated by horizontal lines. Statistical significance was calculated with Student’s unpaired, two-tailed t-test. ***, p=0.0001. (E) Photoactivation of PA-GFP-β-actin in wild-type and twf1/twf2-KO B16-F1 cells co-expressing mCherry-LifeAct as marker for lamel-lipodia, and knockout-rescue cells co-expressing mCherry-TWF-1. Timepoint “Pre” is a frame before photoactivation and 0 s the first frame after the activation. Scale bar = 5 *μ*m. (F) Analysis of the PA-GFP-actin fluorescence decay. Data represent mean of 8-16 experiments with standard deviations shown. Halftimes for GFP-actin fluorescence decays: B16-F1 = 13.1 s, twf-1/2-KO = 22.5 s, twf-1/2-KO + mCherry-TWF-1 rescue = 14.6 s.

We then studied if the decreased F-actin treadmilling in lamellipodia in twf1/twf2-KO cells is linked to defects in F-actin disassembly. For this purpose, we transfected wild-type and twf1/twf2-KO cells with a plasmid expressing photoactivable PA-GFP-β-actin. As a marker for lamellipodial actin filaments, the cells were co-transfected with a plasmid expressing Cherry-LifeAct. By following the decay of PA-GFP-β-actin fluorescence, we detected a significant decrease in the rate of actin filament disassembly at the lamellipodia of twf1/twf2-KO cells, and this could be rescued by expressing mCherry-TWF-1 in the knockout cells (Figure 2E-F, Supplementary movie 3).

Together, the FRAP and photoactivation experiments suggest that twinfilin enhances actin filament disassembly in cells, and hence the twf1/twf2-KO cells display abnormal accumulation of actin filaments at lamellipodia and endocytic structures. The decreased actin filament disassembly rates in the absence of twinfilin are expected to lead to a smaller pool of actin monomers, and consequent decrease in actin filament treadmilling rates as detected in our FRAP experiments on twf1/twf2-KO cells.

### Effects of twinfilin on actin filament barbed end polymerization and depolymerization

Because twf1/twf2-KO cells displayed decreased actin filament disassembly rates compared to wild-type cells, and because previous studies suggested that mammalian twinfilins accelerate actin filament barbed end depolymerization^49^, we analyzed the effect of mouse twinfilin-1 on the dynamics of single actin filaments in vitro^55, 56^ (Figure 3A). With this approach, we detected a similar rate of barbed end depolymerization of bare ADP-actin filaments (~10 subunits/s) as observed in previous studies by bulk fluorometric actin disassembly assay^57^ (Figure 3B). Surprisingly, addition of twinfilin-1 did not accelerate filament barbed end depolymerization as previously reported^48, 49^. Instead, we observed a concentration-dependent decrease in filament barbed end depolymerization rate that plateaued to the level of ~6 subunits/s at saturating twinfilin-1 concentration (Figure 3B). Consistent with earlier single filament TIRF experiments^48, 49^, addition of the N-terminal half of mouse cyclase-associated protein 1 (CAP1) did not enhance the filament barbed end depolymerization in the presence of twinfilin-1 (Figure 3C).

**Figure 3.**
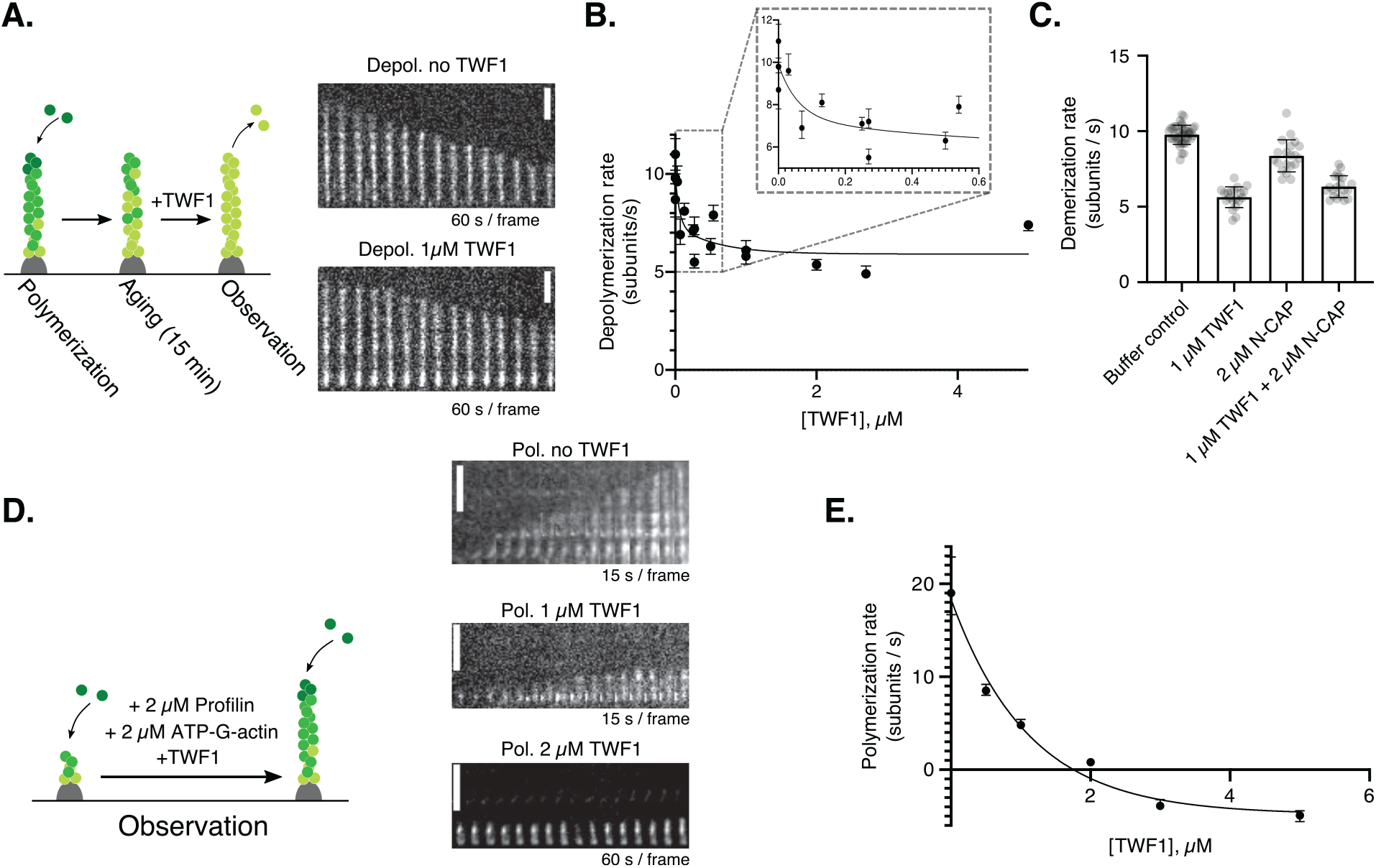
Twinfilin associates with filament barbed ends and does not accelerate depolymerization. (A) Representative examples of individual filament during depolymerization experiment performed in the absence and presence of 1 *μ*M TWF-1. Scale bar = 5 *μ*m. (B) Barbed end depolymerization of ADP-actin filaments in presence of different concentrations of TWF-1. Data points represent median values of at least 20 filaments with quartiles shown. (C) Barbed end depolymerization of ADP-actin filaments in the presence of TWF-1 and/or N-CAP1. Data points represent individual measurements, with mean and standard deviations shown. (D) Polymerization of actin filaments at barbed ends in presence of 2 *μ*M ATP-G-actin and 2 *μ*M profilin, and different concentrations of TWF-1. Scale bar = 5 *μ*m. (E) Analysis of actin filament barbed end polymerization in the presence of 2 *μ*M ATP-G-actin and 2 *μ*M profilin, and different concentrations of TWF-1. Positive and negative values indicate filament polymerization and depolymerization, respectively. Data points represent median values of at least 20 filaments with quartiles shown. Data in panels B and E were fitted with equation Y=(Y_0_ - Y_MIN_) * 10 ^ (−K * X) + Y_MIN_, where Y_0_ is value of Y when X (= concentration of twinfilin) is zero, Y_MIN_ is value of Y in plateau, and K is a rate constant.

We then examined if twinfilin prevents actin filament polymerization at barbed ends, as suggested by earlier fluorometric experiments^47^. For this purpose, the assay was performed under assembly promoting conditions in the presence of 2 *μ*M ATP-G-actin and 2 *μ*M profilin, and by varying the concentration of twinfilin-1 (Figure 3D). Consistent with earlier results^47^, twinfilin-1 inhibited actin filament polymerization at barbed ends. When the twinfilin-1 concentration exceeded the one of profilin/ G-actin complexes, the filaments began to depolymerize (Figure 3E). Collectively, these data suggest that twinfilin sequesters actin monomers and transiently associates with actin filament barbed ends to prevent filament polymerization, but allows their depolymerization with a rate of approximately 6 subunits/s.

### Twinfilin controls the dynamics of CP in cells

The effect of twinfilin on F-actin barbed end dynamics in vitro does not explain why depletion of twinfilins in mammalian cells leads to decreased actin disassembly. Therefore, we focused on its other interaction partners. Twinfilin binds CP through its C-terminal tail^41,42,58^, and both twinfilin and CP localize to lamellipodia in mammalian cells^16,43,59,60^. Whereas CP localizes to the distal edge of lamellipodia^16^, the precise localization pattern of twinfilin has not been reported. We thus expressed EGFP-CP and EGFP-twinfilin-1 in B16-F1 cells and compared their localizations to AlexaFluor-phalloidin labelled actin filaments. Line profile analysis of fluorescence intensity across lamellipodia revealed that whereas CP was enriched at the distal edge of lamellipodial F-actin network, twinfilin localized throughout the lamellipodium, being slightly enriched at the proximal region of the F-actin network (Figure 4A-D). Although twinfilin-1 localized throughout the lamellipodial actin filament network, FRAP experiments on cells expressing EGFP-twinfilin-1 revealed that twinfilin does not display retrograde treadmilling with the lamellipodial actin filament network, but is instead recovers progressively across the whole lamellipodial network, being a dynamic component of the lamellipodium (t_1/2_ = 1.48 s, Figure 4E-F, supplementary movie 4).

**Figure 4.**
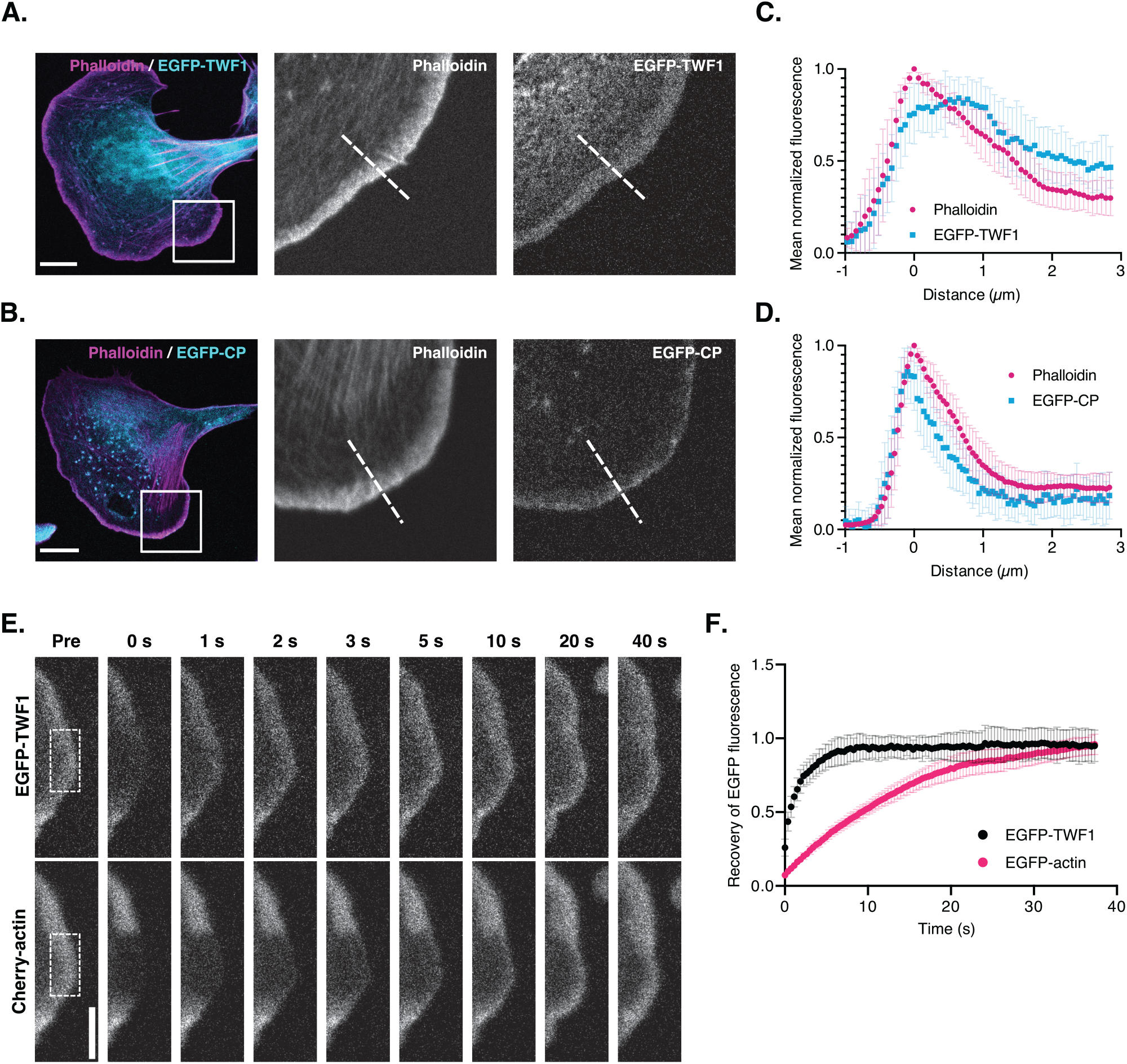
Twinfilin and capping protein display different localizations and dynamics in lamellipodia. (A) Localization of EGFP-TWF-1 in the lamellipodium of a mouse B16-F1 cell. (B) Localization of EGFP-CP in the lamellipodium of mouse B16-F1 cell. Scale bars in panels A and B = 10 *μ*m. Images in the middle and right in panels A and B are magnifications of lamellipodial regions highlighted in the images in left. F-actin was visualized by phalloidin. (C-D) Line profiles generated across the center of lamellipodia as indicated with dotted line. Data represent mean of 22 (panel C) and 24 (panel D) measurements of individual lamellipodia, with standard deviations shown. The ‘0 *μ*m’ value in x-axis is set to correspond the peak intensity of phalloidin. (E) Simultaneous fluorescence recovery after photobleaching of EGFP-TWF-1 and mCherry-actin in a mouse B16-F1 cell lamellipodium. Time-point “Pre” is a frame before bleaching with bleached region indicated with dotted square, and 0 s the first frame after the bleaching. Scale bar = 5 *μ*m. (F) Analysis of EGFP-TWF-1 and EGFP-actin recovery after photobleaching. Data represent mean of 11 and 9 experiments, respectively, with standard deviations shown. Recovery halftimes for EGFP-TWF-1 and EGFP-actin were 1.48 s and 10.19 s, respectively.

We next examined if twinfilin could regulate CP in cells. Expression level of CP was not altered upon depletion of twinfilins (Supplementary figure 1B). However, by comparing the localization of EGFP-CP in wild-type and twf1/twf2-KO cells, we learned that in the absence of twinfilins, CP was no longer restricted to the leading edge of lamellipodium, but its localization instead spread throughout the lamellipodial actin filament network (Figure 5A-C). Moreover, FRAP experiments revealed that also the dynamics of CP was drastically altered in twinfilin-deficient cells. Instead of dynamic exchange at the distal edge of lamellipodium observed in wild-type cells^37^, CP displayed stable association with the actin filament network and underwent retrograde flow in lamellipodia of twf1/twf2-KO cells (Figure 5D, supplementary movie 5). Transient expression of Cherry-twinfilin-1 rescued CP dynamics back to the normal level (Figure 5D-F), confirming that the decrease in CP dynamics in the twf1/ twf2-KO cells was due to lack of twinfilin expression.

**Figure 5.**
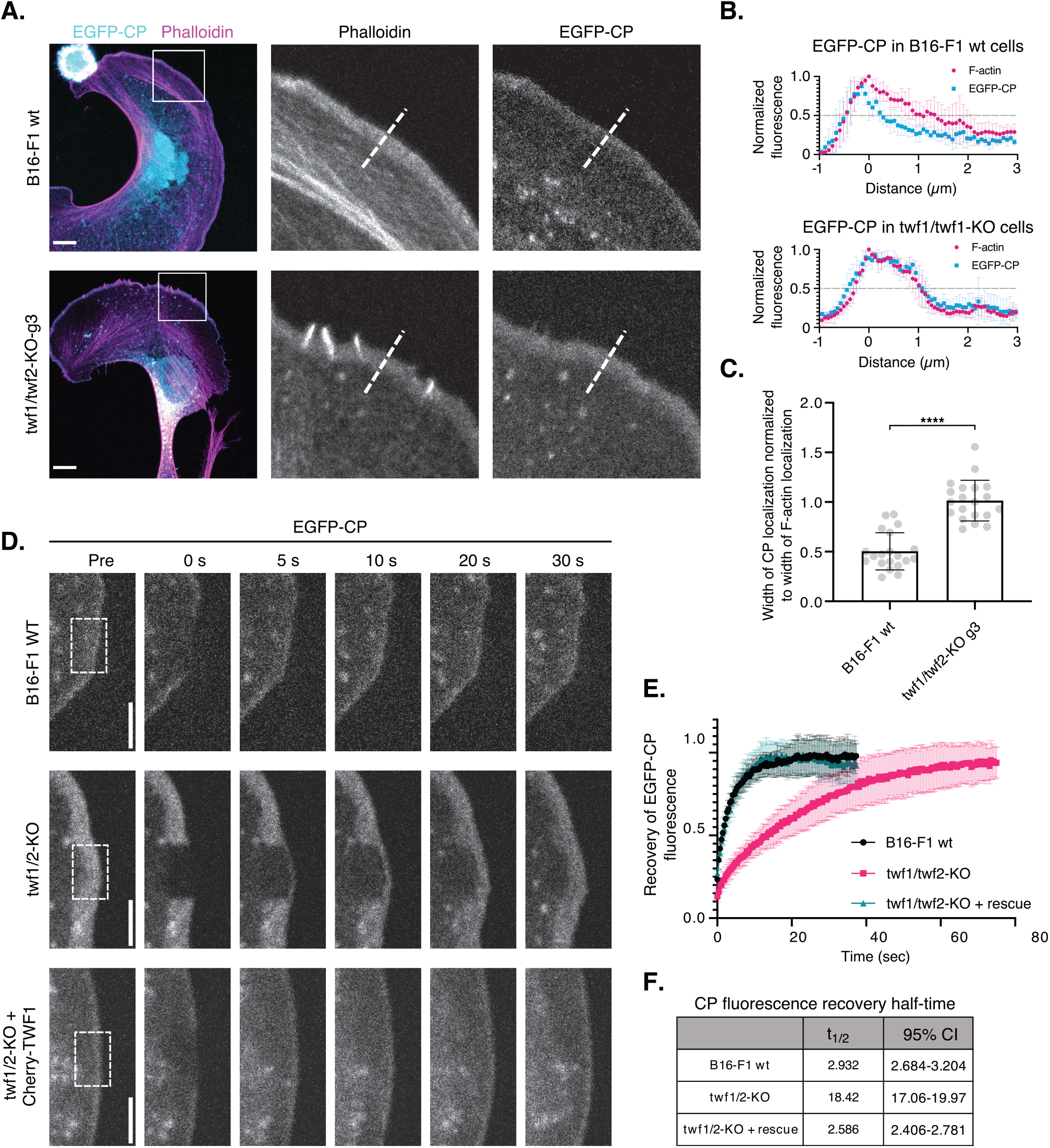
Twinfilin regulates capping protein localization and dynamics. (A) Localization of EGFP-CP in wild-type and twf1/twf2-KO B16-F1 cells, where F-actin was visualized with AlexaFluor-568 phalloidin. Panels in the middle and right are magnifications of lamellipodial regions highlighted in the whole cell images in left. Scale bars = 10 *μ*m. (B) Examples of line profiles generated across the center of lamellipodia as indicated with dotted lines. Data represent mean of 5 measurements of individual lamellipodia, with standard deviations shown. The ‘0 *μ*m’ value in x-axis is set to correspond the peak intensity of phalloidin. (C) The ratio of CP and F-actin co-localization widths were detected by measuring the width of localization at 50% of maximum intensity. Data points represent measurements from individual lamellipodia with mean values and standard deviations shown. Statistical significance was calculated with Student’s unpaired, two-tailed t-test. ****, p<0.0001. (D) Fluorescence recovery after photobleaching of EGFP-CP in wild-type and twinfilin-1/twinfilin-2 knockout, and in knockout cells expressing mCherry-TWF-1. Scale bars = 5 *μ*m. Bleached region in the lamellipodia are highlighted with dotted squares. (E) Analysis of the FRAP experiments. Data represent mean of 9-12 experiments and the standard deviations are shown. (F) Fluorescence recovery halftimes for EGFP-CP in B16-F1 wild-type, twinfilin-1/twinfilin-2 knockout, and EGFP-TWF-1 rescue cells. 95% confidence interval for halftime is shown.

Because both twinfilin and CP localize to endocytic actin filament structures^47, 61^ and twf1/twf2-KO cells exhibited accumulation of F-actin to endosomes, we examined CP dynamics in these structures. Wild-type and twf1/twf2-KO cells were co-transfected with plasmids expressing Cherry-LifeAct to mark actin structures, and EGFP-CP for FRAP analysis (Supplementary figure 6A-B, Supplementary movie 6). These experiments revealed that, similar to lamellipodia, EGFP-CP displayed >10-fold decreased recovery rate in the endosomal actin structures of twf1/twf2-KO cells compared to the wild-type cells (Supplementary figure 6C-D).

Together, these data reveal that twinfilin is responsible for the rapid dynamics of CP in lamellipodia and endocytic structures. Therefore, twinfilin is also critical for CP’s specific localization to the distal edge of lamellipodial actin filament networks in migrating cells.

### Twinfilin uncaps filament barbed ends in vitro

Our cell biological data above suggest that twinfilin may regulate CP dynamics directly by uncapping filament barbed ends. To test this hypothesis, we performed experiments on single filaments inside a microfluidics chamber with purified mouse twinfilin-1 and chicken CP^36^. Actin filaments were polymerized from spectrin-actin seeds bound nonspecifically on the glass surface, and filaments were subsequently capped by CP. In the absence of twinfilin, filament length was very stable indicating that filaments were capped at their barbed ends (Figure 6A). In the presence of twinfilin-1, a significant fraction of filaments began to depolymerize, indicating that they were uncapped (Figure 6B). By performing the assay with different twinfilin concentrations, we estimated a K_D_ around 300 nM of twinfilin to CP-capped actin filament barbed ends and revealed that at saturating conditions twinfilin accelerated filament uncapping by ~6-fold. Importantly, the filament uncapping was further enhanced by including CP-sequestering protein V-1 to the reactions. Whereas V-1 alone does not accelerate CP dissociation, twinfilin and V-1 together increased the uncapping rate by ~40-fold compared to the buffer control (Figure 6C). These in vitro experiments identified twinfilin as an efficient actin filament uncapping factor, which drastically enhances the dissociation of CP from filament barbed ends, especially in the presence of V-1.

**Figure 6.**
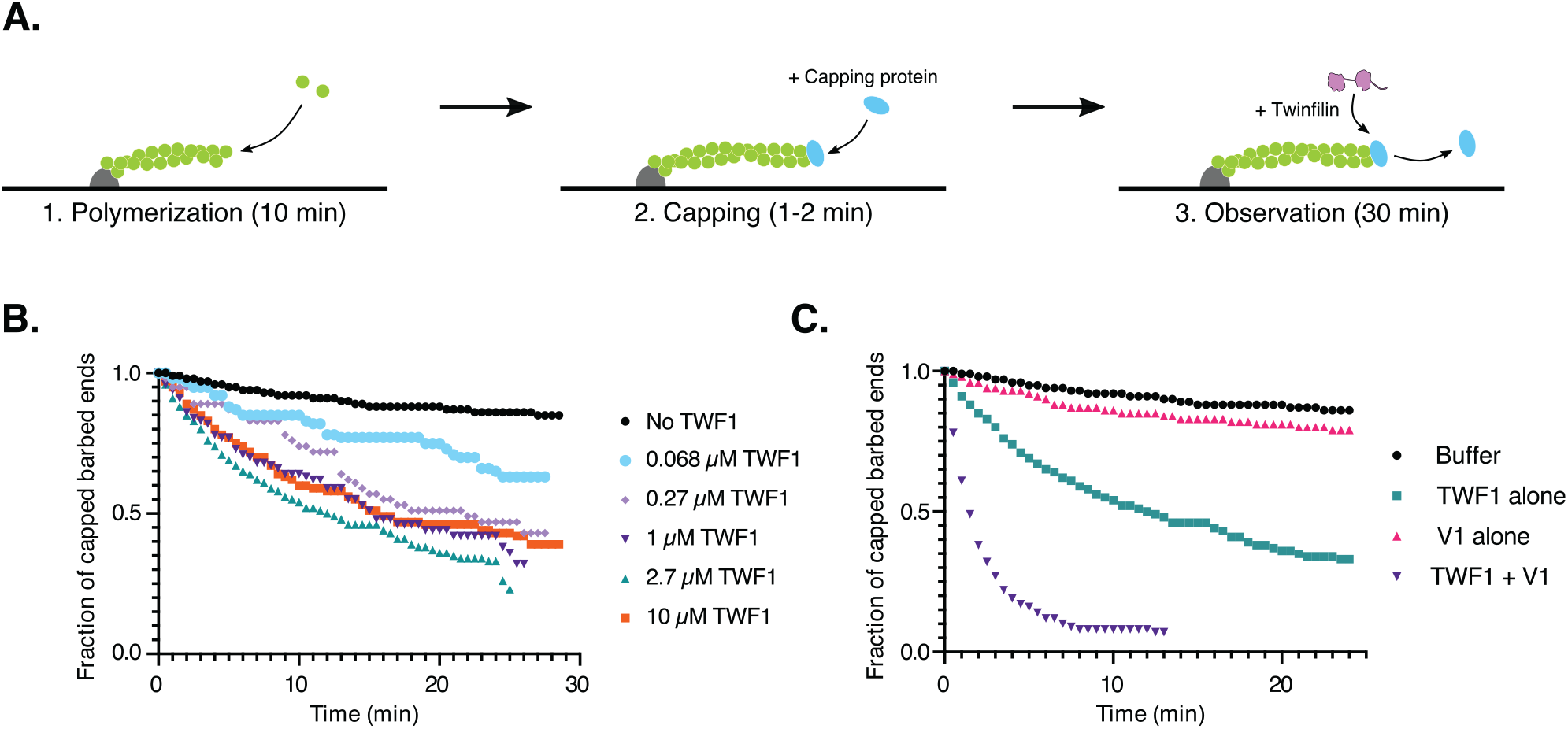
Twinfilin uncaps filament barbed ends in vitro. (A) A schematic overview on the in vitro single filament experimental approach. Actin filaments (10% AlexaFluor-488 labelled G-actin) were polymerized from spectrin-actin seeds and subsequently capped with 100 nM capping protein. Observation was done under constant microfluidics flow of either buffer alone or together with TWF-1 and/or V1 for 60 individual filaments in each condition. (B) Fraction of capped barbed ends over time with different concentrations of TWF-1. (C) Fraction of capped barbed ends over time either without or with 2.7 *μ*M TWF-1 and/or 37 *μ*M V1 protein.

### Actin-binding activity of twinfilin is required for filament uncapping

Twinfilin interacts with CP and binds actin filament barbed ends, and these biochemical functions are dependent of twinfilin’s C-terminal tail and ADF-H domains, respectively^41,42,46,62^. To uncover if twinfilin uncaps filament barbed ends through interacting with CP, actin filament barbed ends, or both, we purified mutant twinfilins that displayed defects in either binding to CP (F323A, K325A, K327A = tail mutant)^42^ or actin (R96A, K98A, R267A, R269A =ADF-H domain mutant)^48, 62^ (Figure 7A). Interestingly, the tail mutant defective in interacting with CP uncapped actin filaments in vitro even more efficiently compared to wild type twinfilin. In contrast, the ADF-H domain mutant did not exhibit detectable uncapping activity, demonstrating that at least in vitro twinfilin’s ability to bind actin filament barbed ends through its two ADF-H domains is critical for dissociation of CP from filament ends (Figure 7B).

**Figure 7.**
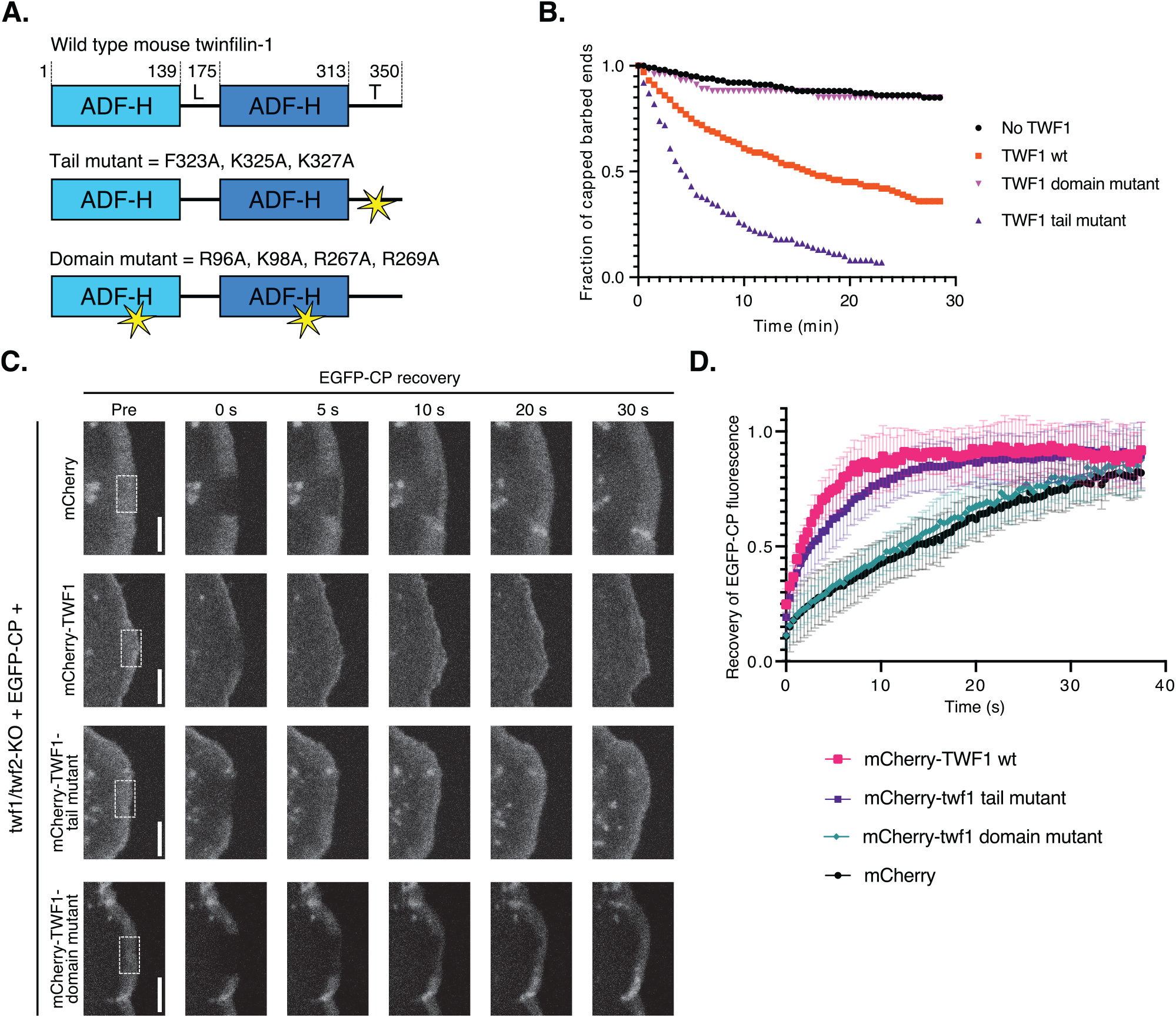
The actin-binding function of twinfilin is required for filament uncapping. (A) Twinfilin-1 mutants used in this study. TWF-1 tail mutant (F323A, K325A, K327A) does not interact with CP, and has decreased affinity for phosphoinositides^42^. TWF-1 domain mutant (R96A, K98A, R267A, R269A) does not interact with actin^48, 62^. (B) Fraction of capped barbed ends over time in 10 *μ*M concentration of wild type or mutant twinfilin-1. Data represents measurements of 60 individual filaments. (C) Fluorescence-recovery-after-photobleaching of EGFP-CP in twf1/twf2-KO cells expressing mCherry (negative control), wild-type mCherry-TWF-1, mCherry-TWF-1 tail mutant, or mCherry-TWF-1 domain mutant. Timepoint “Pre” is a frame before bleaching with region of bleaching indicated with dotted square, and 0 s the first frame after the bleaching. Scale bars = 5 *μ*m. (D) Analysis of fluorescence recovery of EGFP-CP. Data represent mean of 8 measurements for Cherry control, and 12-14 measurements for other constructs, with standard deviations shown. Halftimes of recoveries for EGFP-CP in cells expressing different mCherry-tagged proteins: mCherry-TWF-1 wild-type = 2.4 s, mCherry-TWF-1 tail mutant = 4.2 s, mCherry-TWF-1 domain mutant = 17.1 s, and mCherry (negative control) = 20.4 s.

To examine the effects of the mutant twinfilins on CP dynamics in cells, we expressed mCherry-fusions of these mutants in twf1/twf2-KO cells, and applied FRAP to study if they can rescue the diminished EGFP-CP dynamics (Figure 7C, Supplementary movie 7). Consistent with the in vitro experiments above, the ADF-H domain mutant did not rescue the slow CP dynamics in twf1/twf2-KO cells, whereas the tail mutant rescued the phenotype nearly as well as the wild-type mCherry-twinfilin-1 (Figure 7D). Collectively, the results from in vitro and cell biological experiments demonstrate that twinfilin’s ability to bind actin through its two ADF-H domains is critical for filament uncapping.

## Discussion

Our combined cell biology and single filament imaging studies reveal that the evolutionarily conserved ADF-H domain protein, twinfilin, uncaps actin filament barbed ends to promote rapid CP dynamics and to control the localization of CP within cellular actin filament networks. Moreover, our single filament imaging experiments provide evidence that twinfilin sequesters actin monomers and allows actin filaments to depolymerize following dissociation of CP (Figure 8 A). Consequently, the depletion of twinfilins from cells results in decreased disassembly of actin filament networks.

**Figure 8.**
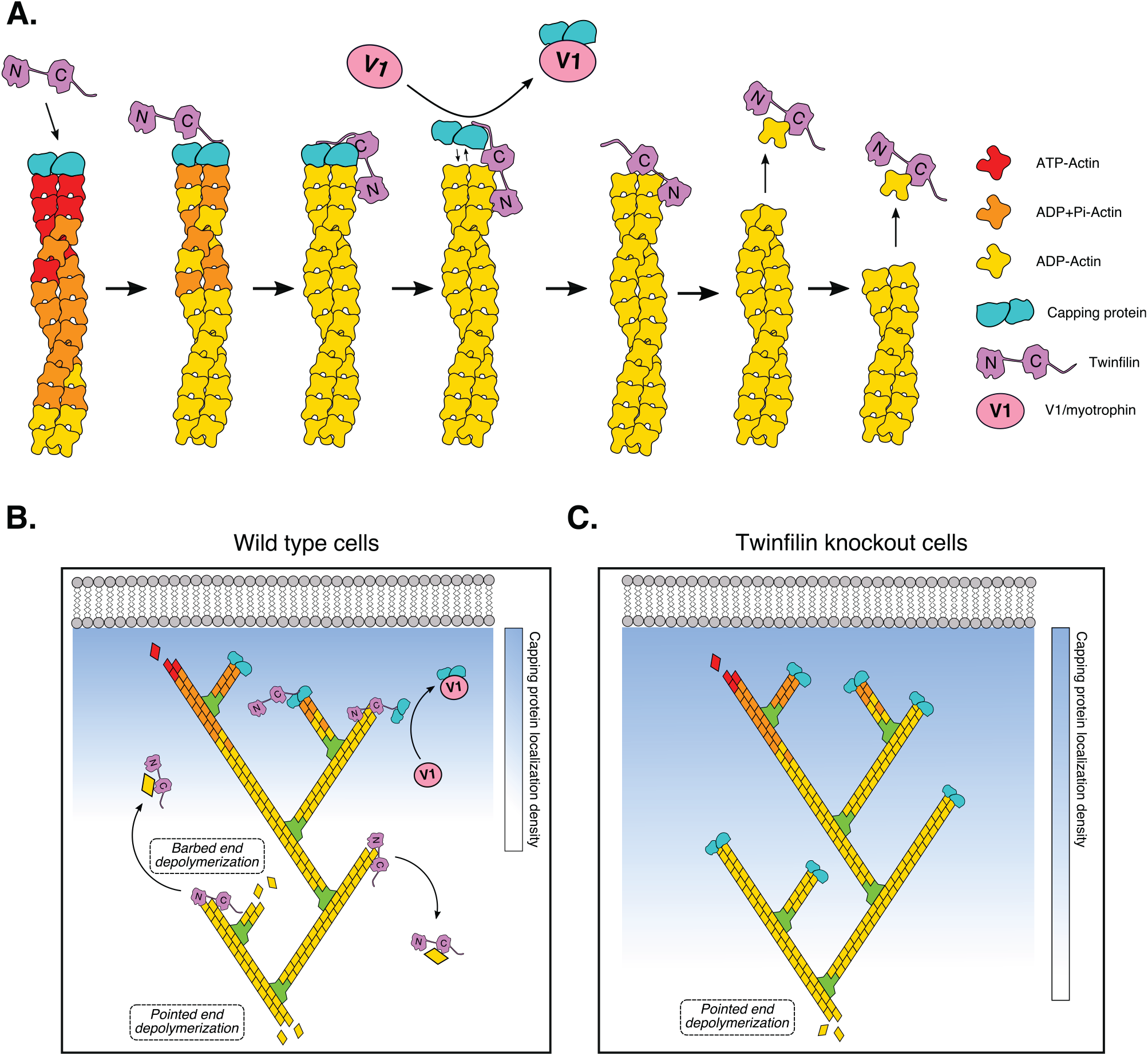
A working model on the role of twinfilin as a regulator capping protein dynamics and actin filament disassembly. (A) Twinfilin dissociates CP from filament barbed end by associating with the filament end through its two ADF-H domains. Additionally, twinfilin binds CP via its C-terminal tail. V1 co-operates with twinfilin to enhance the uncapping rate, most likely by dissociating CP from twinfilin. After uncapping, twinfilin allows dissociation of actin monomers from the barbed ends. (B) In wild-type cells, CP is loaded to barbed ends of lamellipodial actin filaments close to the plasma membrane by CARMIL or other CPI motif proteins. Twinfilin uncaps aged actin filament barbed ends, and thus promotes CP dynamics and restricts its localization to the distal parts of the lamellipodial actin filament network. Aged actin filaments undergo depolymerization from both ends to maintain polymerization-competent pool of ATP-G-actin. (C) In the absence of twinfilin, CP stably caps filament barbed ends throughout the entire lamellipodium. This results in diminished CP dynamics, actin filament barbed end depolymerization, and filaments disassemble only from their pointed ends.

Regulation of CP in cells is orchestrated directly and indirectly by a large number of proteins^9,10,24^. To date, the CPI-motif containing proteins, such as CARMILs, were considered as the best candidates for factors that uncap filament barbed ends, because they decrease the affinity of CP towards barbed ends through allosteric competition^25,26,28,39^. However, to our knowledge there is no direct cell biological evidence to support the role of CARMIL as an uncapping factor. Moreover, CARMIL localizes to the very distal edge of lamellipodia, similar to CP, and CP fails to localize to lamellipodia in CARMIL-depleted cells^31^. Thus, it is more plausible that CARMIL functions as a ‘pro-capping’ protein, which retrieves CP from CP/V-1 complex and allows it to associate with filament barbed ends close to the plasma membrane^9,22,24^. Our cell biological and biochemical work provide strong evidence that twinfilin is the critical factor that is responsible for rapid dynamics and specific subcellular localization of CP in cells.

By using specific twinfilin mutants, we revealed that actin-binding through the two ADF-H domains, but not the interaction with CP, is critical for the actin filament uncapping by twinfilin both in vitro and in cells. Thus, rather than binding to CP through the CPI motif and allosterically enhancing CP-dissociation from barbed ends, twinfilin competes directly with CP for binding to the filament barbed ends and subsequently dissociates CP. Interestingly, the mutant twinfilin unable to interact with CP displayed elevated uncapping activity compared to the wild-type protein. We hypothesize that the CPI-motif in the C-terminal tail of twinfilin can still interact with CP when twinfilin is bound to filament barbed end, thus serving as a transition complex for uncapped/capped barbed ends. This is supported by the fact that the CP-sequestering protein, V-1, enhances uncapping by twinfilin, potentially by dissociating CP from twinfilin. The precise molecular mechanism by which twinfilin interacts with filament barbed ends remains to be elucidated. We note that ADF/cofilins, which are composed of a single ADF-H domain that is structurally similar to the two ADF-H domains of twinfilin^63, 64^, dissociates CP if the ADF/cofilin-decorated segment of filament reaches the barbed end^36^. Therefore, we hypothesize that the first ADF-H domain of twinfilin binds to the side of the filament and changes the conformation terminal actin subunits in filaments, thus weakening the affinity of CP to the barbed ends. Subsequently, the second ADF-H domain of twinfilin associates with terminal actin subunit to displace CP. However, future structural studies are required to reveal the precise mechanism by which twinfilin associates with filament barbed ends.

Earlier study suggested that twinfilin may function as a ‘pro-capping’ factor by loading CP to filament barbed ends by interacting though CPI-motif with CP^43^. It is possible that twinfilin functions both as a CP-activator close to the plasma membrane through its CPI-motif, and as a filament uncapping protein further away from the plasma membrane. However, our FRAP experiments on EGFP-CP are somewhat contradictory with the capping protein activation model^43^. This is because EGFP-CP was efficiently recruited to lamellipodial actin filaments also in the absence of twinfilin, and because inhibition of twinfilin-CP interaction through specific point mutations in the C-terminal tail of twinfilin did not drastically affect CP dynamics in cells. Thus, twinfilin-CP interaction does not appear to be necessary for CP-activation in lamellipodia, suggesting that other CPI motif -containing proteins, such as CARMILs, are the primary activators of CP. However, since functional CARMIL-like proteins have not to our knowledge been reported from fungi^65^, it is possible that interaction with CP through the CPI motif may be more critical for the interplay between twinfilin and CP in yeasts. It is also important to note that an interaction with CP enhances the localization of twinfilin to endocytic actin patches in budding yeast^41, 58^. Thus, another scenario is that the CPI-motif in twinfilin serves as a targeting signal that directs twinfilin to CP-rich actin filaments and enhances filament uncapping activity of twinfilin at least when the cytoplasmic concentration of twinfilin is limiting.

Our biochemical experiments provides evidence that, unlike suggested by earlier studies^48, 49^, twinfilin does not accelerate filament barbed end depolymerization. It is important to note that our experiments were performed on filaments that were attached to coverslips only from their pointed ends (Figure 3A), whereas earlier studies suggesting that twinfilins function as barbed end depolymerizing factors were done on filaments attached to coverslips at multiple points^48, 49^. While the depolymerization rates of bare ADP-actin filaments measured with our approach were similar to the ones obtained with fluorescence-based assays^57^, the depolymerization rates obtained from assays when bare actin filaments were attached to coverslips from multiple points were 5-10 -fold slower^48, 49^. This suggests that attaching actin filaments on coverslips through multiple points may affect their depolymerization rates, and thus provides a plausible explanation for the difference in results between our and earlier studies.

A working a model of how twinfilin regulates the dynamics of lamellipodial actin networks is presented in Figure 8B. Twinfilin restricts the localization of CP to the distal edge of leading edge by uncapping filament barbed ends as filaments display retrograde flow away from the membrane. The trigger for twinfilin to uncap filament barbed ends remains enigmatic at this stage, but it may be related to ATP-hydrolysis in actin. Twinfilin binds ADP-actin filament barbed ends with much higher affinity than ATP-actin barbed ends^47^, and it is possible that twinfilin uncaps filament barbed ends only after nucleotide hydrolysis and Pi-release occurred in the terminal actin subunits of the filament. Our experiments and an earlier study^48^ revealed that at high concentrations twinfilin allows actin filament depolymerization also in the presence of profilin-actin complexes, suggesting that in cells the filaments uncapped by twinfilin can still undergo depolymerization from their barbed ends with a rate of ~6 subunits/s. However, whether twinfilin alone is sufficient to drive actin filament barbed end depolymerization under cellular conditions with a high cytoplasmic concentration of assembly-competent actin monomers, or if it works in concert with other proteins, remains to be elucidated. It is important to note that ADF/ cofilin and cyclase-associated protein can synergistically promote actin filament pointed end depolymerization with a rate up to ~20 subunit/s^66, 67^. Therefore, in wild-type cells the depolymerization of ‘aged’ actin filaments from both barbed and pointed ends may have an important role in actin turnover, as proposed earlier by Wioland et. al.^36^. In the absence of twinfilin, CP remains stably bound to filament barbed ends throughout the lamellipodia, and thus actin dynamics relies entirely on filament depolymerization from their pointed ends. This model also provides an explanation for diminished actin filament disassembly rates in twf1/twf2-KO cells.

In addition to lamellipodial actin dynamics, twinfilin has been linked to endocytosis^51, 68^, lymphoma progression^69^, chemotherapy resistance^52^, cardiac hypertrophy^70^, invasive migration^51, 71^, platelet reactivity and turnover^53^, as well as regulation of cochlear stereocilia length^72^. In the future, it will be important to examine if twinfilin contributes also to these processes by uncapping filament barbed ends, and subsequently allowing filament depolymerization. Moreover, it will be interesting to determine the precise molecular mechanism by which twinfilin, through its ADF-H domains, dissociates CP from the filament barbed end.

## Materials and methods

### Antibodies and reagents

Rabbit anti-mouse twinfilin-1 antibody [dilution in Western blot (WB), 1:500] was described earlier^45^. Other antibodies used in the study were: Rabbit anti-twinfilin-2 antibody (Sigma-Aldrich #HPA053874, WB, 1:100), rabbit anti-CAPZß antibody (Sigma-Aldrich, #HPA031531, WB, 1:100), mouse anti-α-tubulin antibody (Sigma-Aldrich, #T5168, WB 1:10,000), mouse anti-ß-actin antibody (Sigma-Aldrich, #A5441, WB, 1:10,000), Rabbit anti-p34-Arc/ARPC2 (Merck Millipore, #07-227, dilution in immunofluorescence (IF), 1:200), goat anti-Rabbit IgG AlexaFluor-488 conjugated secondary antibody (Thermo Fisher, #A-11034, IF, 1:400), goat anti-Rabbit IgG AlexaFluor-568 conjugated secondary antibody (Thermo Fisher, #A-11011, IF, 1:400), goat anti-Rabbit IgG AlexaFluor-647 conjugated secondary antibody (Thermo Fisher, #A-32733, IF, 1:400), goat anti-Mouse IgG HRP conjugated secondary antibody (Thermo Fisher, #31430, WB, 1:10,000), goat anti-Rabbit IgG HRP conjugated secondary antibody (Thermo Fisher, #32460, WB, 1:1,000). Other reagents used in the study were: CellMask Deep red (Thermo Fisher, #H32717, IF, 1:25,000), AlexaFluor-647-transferrin (Thermo Fisher, #T23366, IF, 1:400), AlexaFluor-488-phalloidin (Thermo Fisher, #A12379, IF, 1:400), AlexaFluor-555-phalloidin (Thermo Fisher, #A34055, IF, 1:400), AlexaFluor-568-phalloidin (Thermo Fisher, #A12380, IF, 1:400), AlexaFluor-647-phalloidin (Thermo Fisher, #22287, IF, 1:400), and DAPI (Thermo Fisher, #D1306).

### Plasmids

Mouse twinfilin-1 cDNA was cloned to pEGFP-C1A vector by using the SpeI/HindIII cloning sites, to pmCherry-C1 vector with SacI/HindIII, and to pHAT vector with NcoI/HindIII. Point mutations to twinfilin-1 cDNA were introduced using quick-change approach. EGFP-CP-ß2 was a gift from Dorothy Schafer (University of Virginia, Chalottesville, USA)^60^, EGFP-ß-actin and mCherry-ß-actin were gifts from Martin Bähler (Westfalian Wilhelms-University Münster, Germany), pSpCas9(BB)-2A-GFP a gift from Feng Zhang (Addgene plasmid #48138)^73^, mPA-GFP-actin was a gift from Michael Davidson (Addgene plasmid #57121), mCherry-LifeAct was a gift from Maria Vartiainen (University of Helsinki, Finland), and pGST-Tev-V1 a gift from John A. Hammer (NIH National Heart, Lung and Blood Institute, USA). pET-3d plasmid containing Chicken Capping Protein α1 and β1 subunits was a gift from John Cooper (Addgene plasmid #13451) The N-terminal domain of mouse CAP1 in pSUMOck4 was described earlier^66^.

### Cell lines

Mouse B16-F1 cell line was purchased from Sigma-Aldrich (#92101203, LOT #12F003). Cells were cultured in DMEM (Lonza, #BE12-614F) supplemented with 10% FBS (Gibco, #10500-064, LOT#08Q2061K and LOT#2025814K) and penicillin-streptomycin-glutamine solution (Gibco #10378016). Cells were frequently tested against mycoplasma with MycoAlert mycoplasma detection kit (Lonza #LT07-418). Cells were transfected with Fugene HD transfection reagent (Promega #E2312). For live cell imaging, culture media was changed to either phenol red -free DMEM (Gibco, #21063029) or DMEM^EGFP-2^ anti-bleaching live cell visualization medium (Evrogen, #MCK02), both supplemented with 10% FBS and GlutaMAX-I (Gibco, #35050061).

### CRISPR-Cas9

Twinfilin knockout cell lines were generated with non-homologous end-joining approach as described earlier^73^. Shortly, target sequences for twinfilin-1 exon 3 (guide 1: GGAGTCTGAAGGTGGACTAC, guide 2: ACGGCTGCTTGTCCTCCAGC), twinfilin-2 exon 3 (guide 3: GCACAGCCCGGTCGTAGTCC) and twinfilin-2 exon 4 (guide 4: CTTCTTCACGGTGGCGCGTG) were cloned into pSpCas9(BB)-2A-GFP vector as described earlier^73^. B16-F1 cells were transfected with the above-mentioned plasmids and GFP-positive cells were sorted with BD FACSaria II cell sorter. Twinfilin knockout cell lines were identified with Western blot with TWF-1 and TWF-2 specific antibodies. The genomic DNAs from wild-type B16-F1 and twinfilin knockout cells were extracted with Genomic DNA extraction kit (Invitrogen, #K1820-00) and exon 3 regions were sequenced with twinfilin-1 specific primers 5’-AAGACTGCCGCTTCTAACCC and 5’-GAGTTGAGACCTACGTCACTC, and with twinfilin-2 specific primers 5’-GAGGAGAATGTGGGATGTGCC and 5’-CTCGTCTGTTCTCCCCACTT at Eurofins Genomics sequencing service (for Sanger sequencing) and Institute of Biotechnology Sequencing unit at University of Helsinki (for NGS MiSeq sequencing)^75^. Sequence analysis was performed with Geneious 6.1.8 (Biomatters Ltd.)

### Western blot

Cells were rinsed with cold PBS and lysed with lysis buffer [0.1% Tritox X-100, 4 mM EDTA, cOmplete ultra EDTA-free protease inhibitor cocktail (Roche Applied Science, #06538282001) in PBS] and sonicated thoroughly. Cell lysates were cleared with centrifugation and protein concentrations were measured with Protein Assay kit (Bio-Rad, #5000001). 25-50 *μ*g protein were loaded to 10-well SDS-PAGE gradient gel (Bio-Rad 456-1094). Proteins were transferred to nitrocellulose membrane (Bio-Rad, #170-4158/#170-4159), which were blocked with 5% milk in TBS-T (0.05% Tween-20). Primary antibodies were incubated overnight at +4°C in 5% milk in TBS-T or 5% BSA in TBS-T. Membranes were washed with TBS-T, and incubated with secondary antibody in 5% BSA in TBS-T. After washing with TBS-T, detection of proteins from the membranes was performed with Western Lightning ECL Pro solution (Perkin Elmer, #NEL121001EA).

### Immunofluorecence stainings

Cells were cultured on 20 *μ*g/ml laminin (Sigma-Aldrich, #L2020) -coated cover slips or CellCarrier Ultra 96 well plates (Perkin Elmer, #6055302). Cells were fixed with 4% PFA in PBS, washed several times with PBS, permeabilized with 0.05% Triton X-100 in PBS, and washed again with PBS. Cells were blocked with 1% BSA in PBS for one hour at room temperature and incubated with primary antibody in 1% BSA/PBS for two hours at room temperature. Cells were washed several times with 0.2% BSA in Dulbecco’s PBS, and incubated with secondary antibody, AlexaFluor-conjugated phalloidin, CellMask and DAPI in 1% BSA/PBS. Cells were mounted to microscopes slides with Vectashield Vibrance antifade mounting media (Vector Laboratories, #H-1700).

### Confocal imaging

Imaging was performed using Leica TCS SP8 STED 3X CW 3D confocal microscope with HC PL APO 93x/1.30 motCORR STED white objective or Leica TCS SP8 X white light laser confocal microscope with 63x HC PL Apo CS2 objective. Maximum projection images from stacks were generated and the images were analyzed with Fiji/ImageJ. For line profile analysis, line width of 5 pixels was used, and lines were drawn across the center of the lamellipodium.

### High-content imaging

Imaging was performed with Opera Phenix High content screening system (Perkin Elmer) with 40x NA 1.1 water immersion objective. Maximum projections from stack images and illumination correction were performed with CellProfiler 3.1.8 (www.cellprofiler.org). Cells were segmented with CellProfiler based on DAPI and CellMask fluorescence, and the Arp2/3-complex positive actin structures and endosomal actin structures based on p34-antibody staining and AlexaFluor-transferrin fluorescence, respectively. Data were analyzed with Prism 8 (Graphpad inc.).

### Lamellipodia protrusion assay

Cells were cultured on CellView 35 mm glass-bottom cell culture dishes (Greiner Bio-One, #627861) coated with 20 *μ*g/ml laminin for one hour prior to imaging. DIC imaging was performed with 3i Marianas (3i intelligent Imaging Innovations) with 63x/1.2 W C-Apochromat Corr WD=0.28 M27 and DIC slider. Kymographs of protrusive lamellipodia were generated and lamellipodia protrusion velocities were measured with Fiji/ImageJ.

### Fluorescence recovery and photoactivation assays

For fluorescence recovery after photobleaching and fluorescence decay after photoactivation assays, cells were transfected as described above. After 24 hours, cells were transferred to CellView 35 mm glass-bottom cell culture dishes (Greiner Bio-One, #627861) coated with 20 *μ*g/ml laminin. Media was exchanged to DMEM^GFP-2^ anti-bleaching live cell visualization medium (Evrogen # MCK02) supplemented with 10% FBS and GlutaMAX (Gibco #35050061) and 20 *μ*g/ml rutin. Imaging was performed with Leica TCS SP5 II HCS-A confocal microscope with HCX PL APO 63x/1.2 w objective and FRAP booster. Acquisition was done with Leica LAS AF 2.8.8 software and image analysis with either Leica LAS AF 2.8.8 or Fiji/ImageJ as described earlier^74^. Briefly, three frames were imaged with low laser power (488/35mV with 1-5% laser power) and rectangular region was subsequently bleached with 5-7 iterations at full laser power. In photoactivation, three iterations of UV laser (405 nm/50 mW) was used at full laser power. Fluorescence recovery and decay were then followed with identical setting to pre-bleach/activation frames. Fluorescence intensity was measured from identically sized regions from the bleached (or photoactivated in case of photoactivation experiments) region, from a cytoplasmic region (bleaching control), and from a region outside (background) the cell. After background subtraction and correction of fluorescence bleaching during imaging, recovery (or fluorescence decay in case of photoactivation experiments) rate was normalized to the pre-bleach intensity. Data were analyzed with Microsoft Excel, and recovery as well as fluorescence decay curves were fitted to calculate the recovery halftimes with Prism 8 (Graphpad Inc.). Data were fitted with the equation F=(F_0_ − F_MIN_)*10^(−K*t) + F_MIN_, where F_0_ is fluorescence in t = 0 s, F_MIN_ is fluorescence in plateau, K is rate constant, and t is time. t_1/2_ = ln(2)/K.

### Protein purification

His-tagged recombinant twinfilins were expressed in E.coli BL21 (DE3) pLysS cells in autoinduction Luria broth (Formedium) overnight at +18°C. Cells were harvested with centrifugation, suspended in lysis buffer (20 mM Tris-HCl, pH 7.5, 150 mM NaCl, 25 mM imidazole), and sonicated thoroughly. Lysate was cleared with centrifugation and loaded to HisTrap FF crude 5 ml Ni-NTA column (GE healthcare, #17-5286-01). Column was washed with 10 column volume of lysis buffer. His-TWF-1 was eluted with elution buffer (20 mM Tris-HCl, pH 7.5, 150 mM NaCl, 250 mM imidazole). His-TWF-1 was further purified with Superdex-75 increase 10/300 GL gel filtration column (GE Healthcare) equilibrated 20 mM Tris-HCl, pH 7.5, 100 mM NaCl buffer. Proteins were concentrated with Amicon Ultra-15 Ultracel 10k centrifugal filter units (Merck Millipore, #UFC901024). Purification of cytoplasmic actin, spectrin-actin seeds and recombinant human profilin-1 were performed as described^36^. Recombinant N-terminal half of CAP1 (N-CAP1) and chicken capping protein α1/β2 were expressed and purified as described in earlier study^66^. GST-V-1 recombinant protein was expressed in E.coli BL21 (DE3) pLysS cells) overnight at +18°C. Cells were resuspended to lysis buffer (20 mM Tris-HCl, pH 7.5, 150 mM NaCl) and sonicated thoroughly. Lysate was cleared with centrifugation and bound to Glutathione sepharose 4 Fast Flow beads (GE Healthcare, #17-5132-02). Beads were washed several times with lysis buffer and protein was eluted from beads with 10 mM reduced glutathione in lysis buffer. Elute was further purified with Superdex-75 increase 10/300 GL gel filtration column as described above.

### Single filament imaging experiments

Actin was fluorescently labelled (10% AlexaFluor-488 labelled actin) and single filament imaging setup with microfluidics was described earlier^36^. For barbed end depolymerization assay, equal amounts of ATP-G-actin and profilin were used for polymerizing F-actin from spectrin-actin seeds bound to glass surface in F-buffer (5 mM Tris-HCl pH 7.8, 50 mM KCl, 1 mM MgCl_2_, 0.2 mM EGTA, 0.2 mM ATP, 10 mM DTT and 1 mM DABCO). Filaments were then aged for ~15 min in F-buffer supplemented with 0.1 *μ*M ATP-G-actin to get most of monomers in filament in ADP-form. Different concentrations of TWF-1 and/or N-CAP were injected to the system in F-buffer with microfluidics. Filaments were imaged with Nikon TiE or TE2000 inverted microscopes and depolymerization rate measured with ImageJ or homemade Python algorithm as described earlier^36^. In polymerization assay, 400 nM ATP-G-actin or 2 *μ*M ATP-G-actin supplemented with 2 *μ*M profilin-1 was injected to system with microfluidics along with different concentrations of TWF-1. Filament polymerization rate was measured as above. To measure F-actin uncapping, F-actin (10% AlexaFluor-labelled actin) was polymerized from spectrin-actin seeds as described above and then capped with 100 nM CP. Different concentrations of TWF-1 and V1 proteins were then injected into the system with microfluidics. Uncapping was measured by following when filaments start to depolymerize as described earlier^36^.

### Statistics

All statistical analyses were performed with Prism 8 (Graphpad Inc.). All box-and-a-whisker plots contain median value, the 25th and the 75th percentiles, and the smallest and the largest values. The normality of data was tested with Prism 8. Statistical tests for normally distributed data were done with non-paired two-tailed Student’s t-test, and for non-normally distributed data with two-tailed Mann-Whitney rank sum test.

## Acknowledgements

We thank Tommi Kotila, Ville Paavilainen, and Juha Saarikangas for critical reading of the manuscript. Institute of Biotechnology Light Microscopy Unit, Biomedicum Imaging Unit and Institute of Molecular Medicine Finland High Content Imaging and Analysis Unit are acknowledged for providing support in imaging and image analysis. We thank Mirva Tirkkonen for technical assistance. This study was supported by grants from Academy of Finland (302161) and Cancer Society Finland (4705949) to P.L., from Doctoral School in Health Sciences at University of Helsinki to M.H., and from the Agence Nationale de la Recherche (Grant Muscactin) to G.R-L. and from the European Research Council (Grant StG-679116) to A.J.

## Author contributions

P.L. and M.H. crafted the original idea, and M.H., H.W., A.J., G.R-L., and P.L. designed the experiments. M.H., M.T., and H.W. performed the experiments, and M.H. and H.W. analyzed the data. M.H. and P.L. drafted the manuscript with contribution from all authors. P.L., A.J., and G.R-L. acquired the funding.

## SUPPLEMENTARY FIGURES

**Supplementary figure 1.**
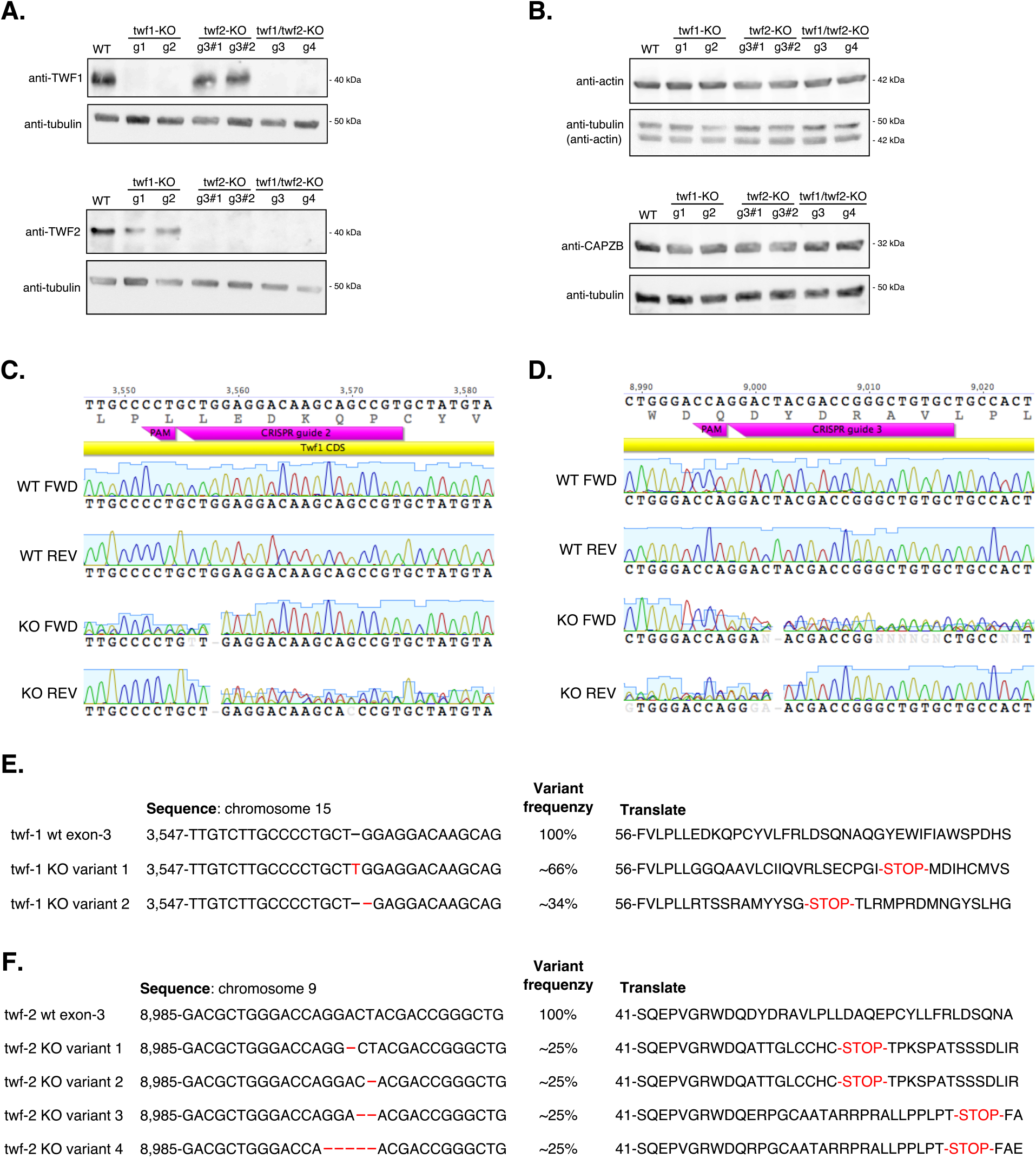
Generation of twinfilin-1, twinfilin-2, and twinfilin-1/twinfilin-2 knockout B16-F1 cells. **(A)** Western blot analysis of wild-type and twinfilin knockout cells performed with anti-TWF-1 and anti-TWF-2 antibodies. Anti-α-tubulin antibody was used as a loading control. **(B)** Western blot analysis of wild type and twinfilin knockout cells for β-actin and capping protein levels. Anti-α-tubulin antibody was used as a loading control. We note that anti-β-actin antibody remained in the blot even after vigorous stripping of the membrane and was visible in anti-α-tubulin detection **(C)** Sanger sequencing of the third exon of *twinfilin-1* gene from genomic DNA of wild-type and twf1/twf2-KO g3 knockout cell lines. Sequences obtained with forward and reverse primers for the third exon of *twinfilin-1* are shown. **(D)** Sanger sequencing of the third exon of *twinfilin-2* from genomic DNA of wild-type and twf1/twf2-KO g3 knockout cell lines. Sequences with forward and reverse primers for the third exon of *twinfilin-2* are shown. **(E-F)** MiSeq NGS sequencing results of *twinfilin-1* exon 3 **(E)**, and *twinfilin-2* exon 3 **(F)** of wild-type and twf1/twf2-KO-g3 cells. Due to chromosome amplification, *twinfilin-1* knockout was a result of two allele variants and *twinfilin-2* knockout a result of four allele variants. All mutant alleles lead to premature stop-codons in the region encoding twinfilins’ N-terminal ADF-H domain. Numbering of nucleotide and amino acid sequences represent the first nucleotide and amino acid residues shown, respectively.

**Supplementary figure 2.**
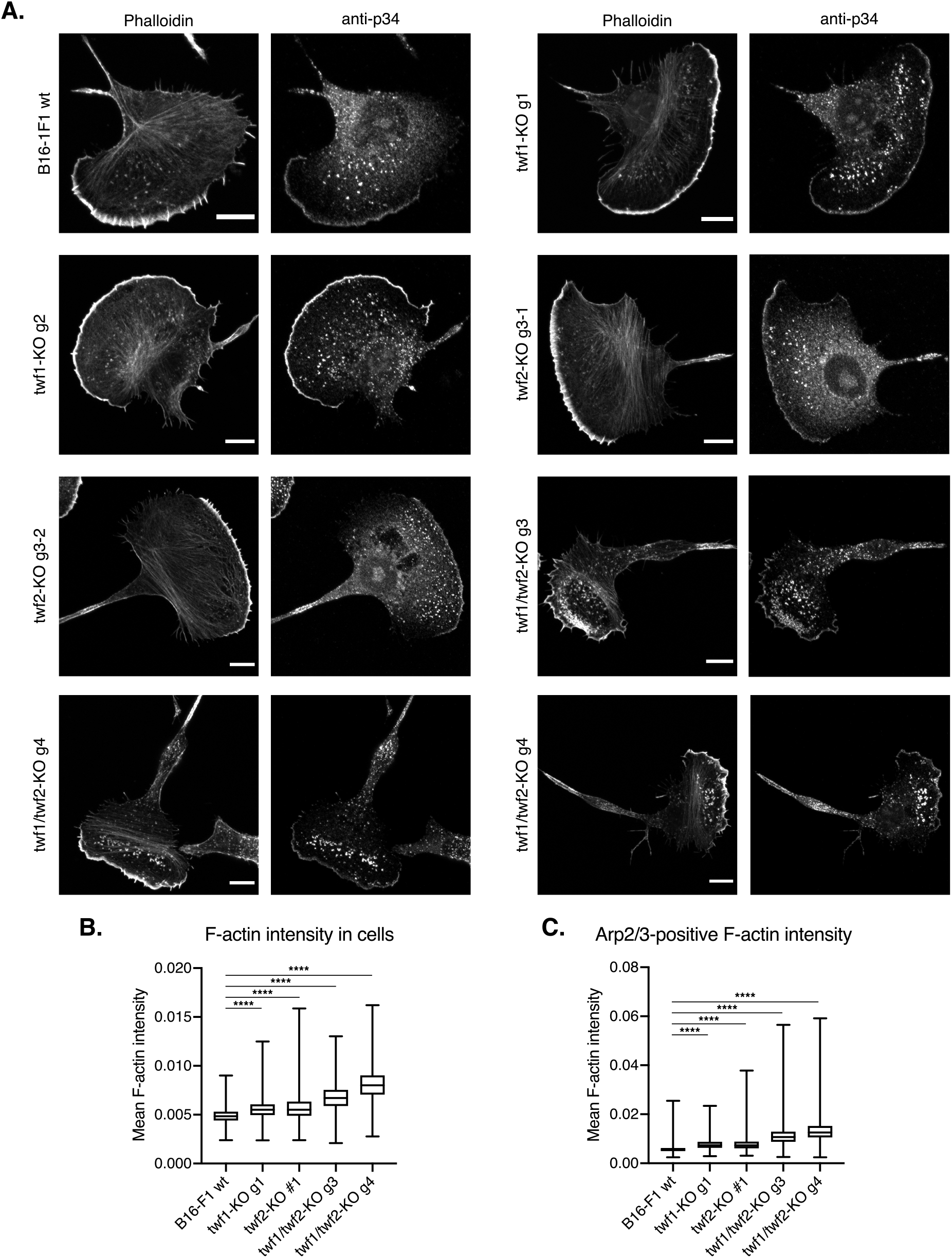
Representative images of wild-type and twinfilin knockout B16-F1 cells. **(A)** B16-F1 cells were stained with AlexaFluor-568 phalloidin (F-actin) and anti-p34 antibody (the Arp2/3 complex). Scale bars = 10 *μ*M. **(B)** Mean F-actin intensity in B16-F1 wild-type and twinfilin knockout cells. Number of measured cells were: B16-F1 wt = 1,958, twf1-KO-g1 = 1,045, twf2-KO-g3#1 = 1,285, twf1/2-KO-g3 = 1,265, twf1/2-KO-g4 = 1,707. **(C)** Mean F-actin intensity in the Arp2/3 complex positive regions of B16-F1 wild-type and knockout cells. The Arp2/3 complex positive regions were identified based on p34-antibody staining. Number of measured cells were: B16-F1 wt = 1,658, twf1-KO-g1 = 884, twf2-KO-g3#1 = 1,009, twf1/2-KO-g3 = 1,100, twf1/2-KO-g4 = 1157. Statistical significances in panels B and C were calculated with Mann-Whitney two-tailed test. ****, p<0.0001.

**Supplementary figure 3.**
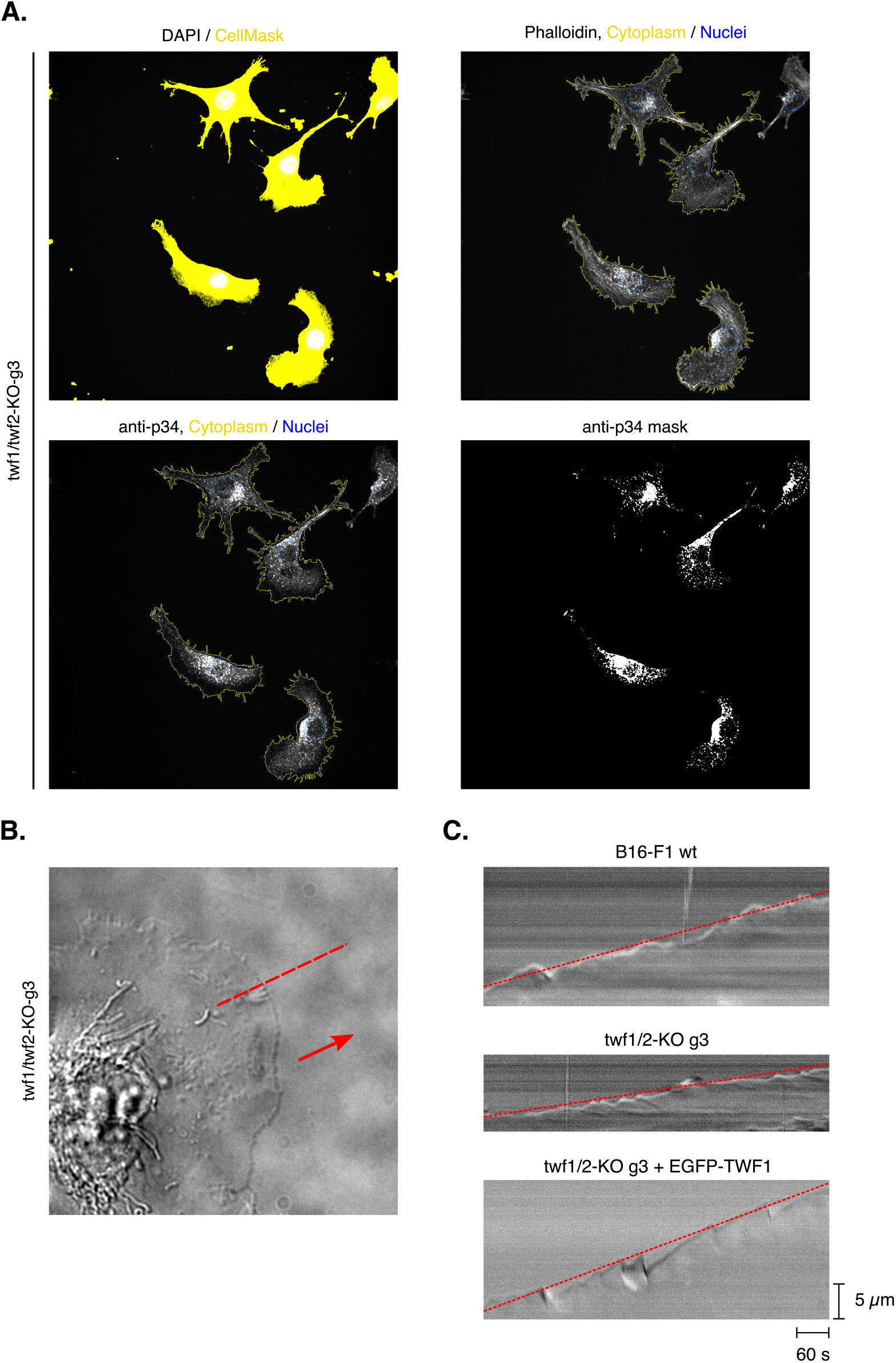
Examples of high-content and lamellipodia protrusion analysis. **(A)** Representative images of segmentation procedure in high-content image analysis. Wild-type and twinfilin-deficient B16-F1 cells were stained with DAPI and CellMask Deep Red to segment nuclei (outlined with blue) and cytoplasm (outlined with yellow). The Arp2/3-complex positive structures were detected with anti-p34 antibody staining and used as a mask for the Arp2/3-complex positive F-actin structures (the most-right panel). F-actin was stained with AlexaFluor-568 phalloidin. Cells touching the border of images were excluded from analysis. **(B)** A representative example of twf1/twf2-KO cell migrating on laminin coated glass imaged with DIC. Direction of migration is indicated with an arrow and kymographs were generated with line drawn across the lamellipodium as indicated with dotted red line. **(C)** Representative examples of kymographs generated from DIC time-lapse images of wild-type, twf1/twf2 knockout, and EGFP-TWF-1 rescue cells. Protrusions velocities were measured from the overall cell front protrusion as indicated with dotted red lines.

**Supplementary figure 4.**
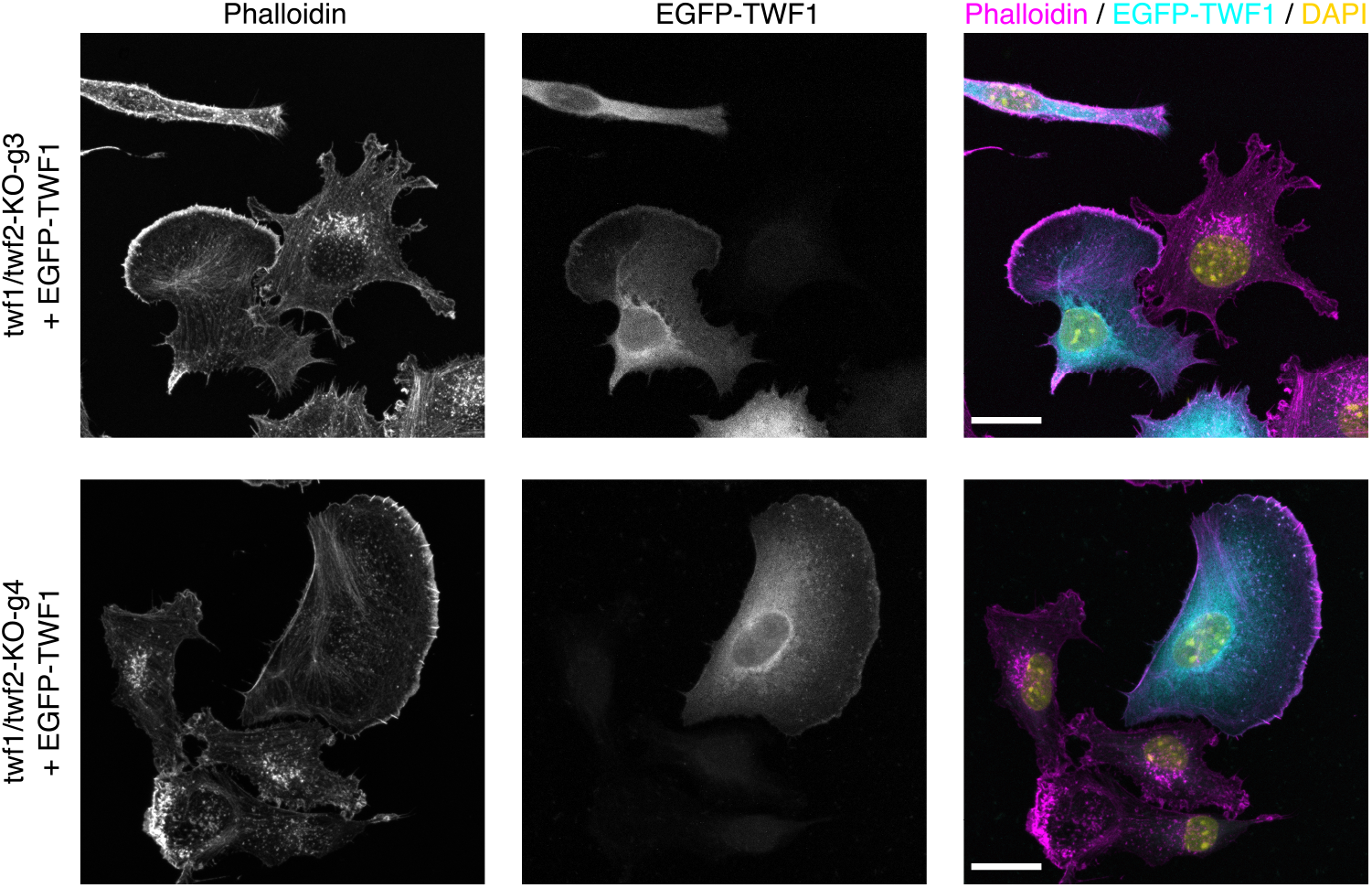
Expression of EGFP-TWF-1 rescues the twf1/twf2 knockout phenotype. Representative images of twf1/twf2-KO cells expressing EGFP-TWF-1 and stained with AlexaFluor-647 phalloidin and DAPI to visualize F-actin and nuclei, respectively. Scale bars = 20 *μ*m.

**Supplementary figure 5.**
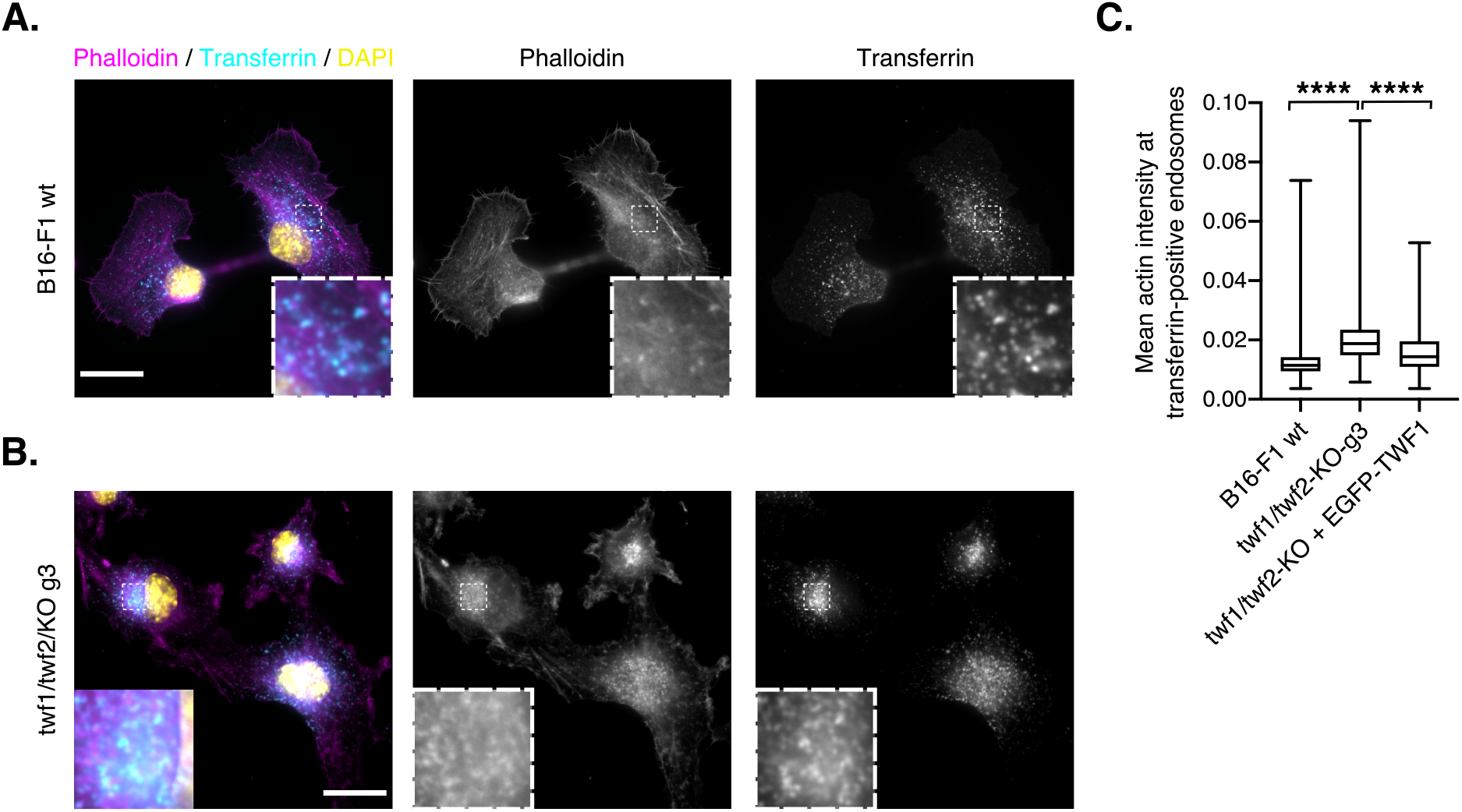
Twinfilin-1/twinfilin-2 knockout leads to an accumulation of F-actin on endosomes at the perinuclear region. **(A)** Phalloidin staining of wild-type B16-F1, and **(B)** twf1/twf2-KO cells after 7.5 min uptake of 20 *μ*g/ml AlexaFluor-647 transferrin. Scale bar = 20 *μ*m. The dotted square indicates the perinuclear region magnified in the insert. **(C)** Mean F-actin intensities in transferrin-positive endosomes of wild-type and twf1/twf2-KO cells as measured from AlexaFluor-555 phalloidin stained cells after 7.5 min intake of 20 *μ*g/ml AlexaFluor-647 phalloidin. Numbers of measured cells were: B16-F1 wt = 2,518, twf1/2-KO-g3 = 3,637, twf1/2-KO-g3 + EGFP-TWF1 = 158. Statistical significances were calculated with Mann-Whitney two-tailed test. ****, p<0.0001.

**Supplementary figure 6.**
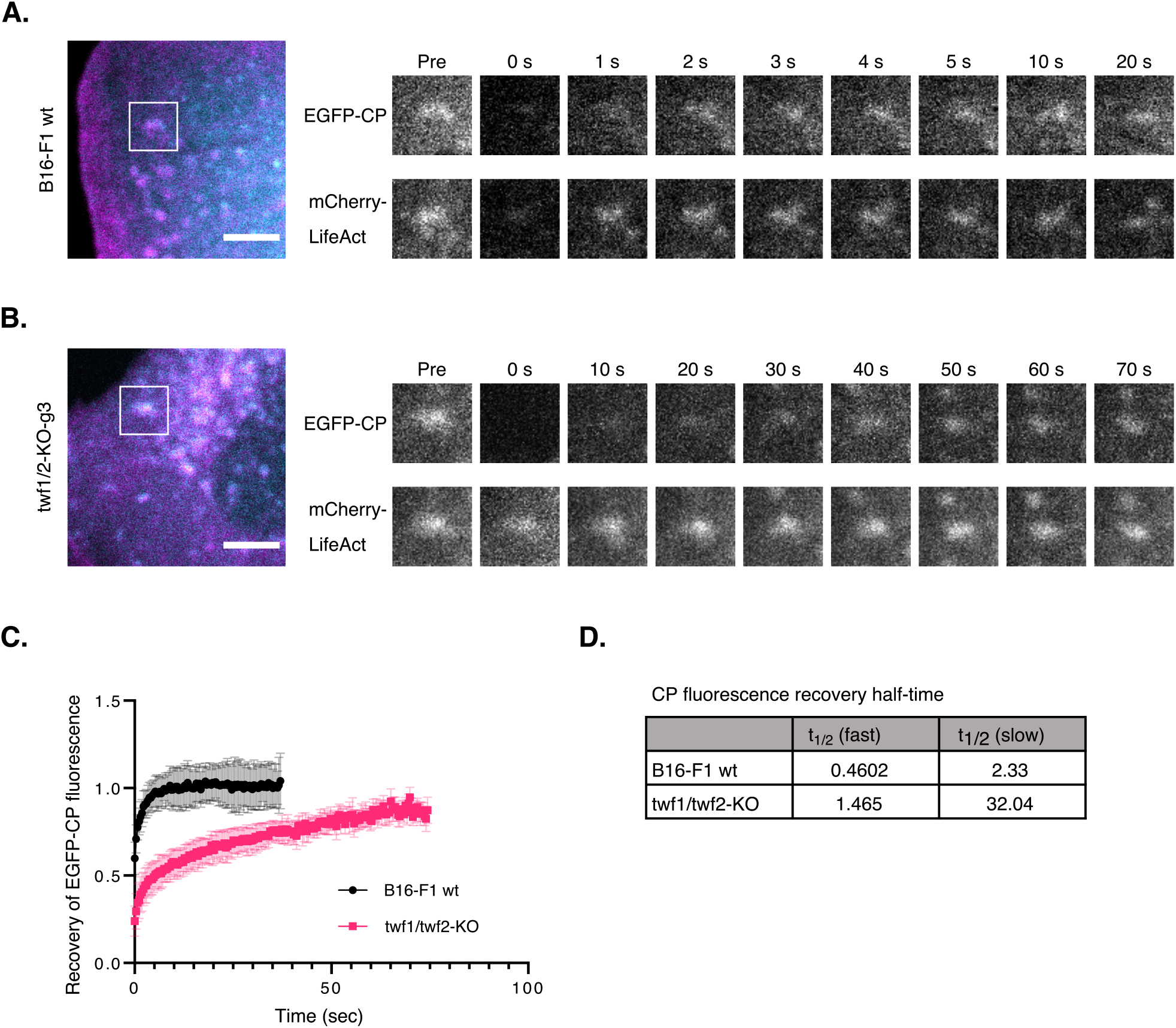
Knockout of twinfilins leads to diminished capping protein dynamics on endosomal actin structures. **(A)** Fluorescence recovery after photobleaching experiment on wild-type, and **(B)** twf1/twf2-KO cells expressing EGFP-CP (cyan) and mCherry-LifeAct (magenta). The recovery of EGFP-CP fluorescence was measured from LifeAct-positive EGFP-CP puncta. Scale bars = 10 *μ*m. Regions of interest are highlighted with white squares and magnified in panels on right with indicated time points. **(C)** Recovery of EGFP-CP fluorescence over time in wild-type and twf1/twf2-KO cells. Each data point represent mean of 8 and 10 individual experiments, respectively, and standard deviations are shown. **(D)** Halftimes of EGFP-CP fluorescence recoveries. The half-times for both fast (diffuse fraction) and slow (bound fraction) are shown.

## SUPPLEMENTARY MOVIE LEGENDS

**Supplementary movie 1**. Lamellipodia protrusion of wild-type (left) and twf1/twf2-KO (right) B16-F1 cells on laminin-coated glass-bottom dishes.

**Supplementary movie 2**. Fluorescence recovery of EGFP-actin after photobleaching in wild-type (left) and twf1/twf2-KO (right) B16-F1 cells.

**Supplementary movie 3.** Fluorescence decay of PA-GFP-actin (left panel) after photoactivation in wild-type (upper panel) and twf1/twf2-KO (middle panel) B16-F1 cells, and in twf1/twf2-KO cells expressing mCherry-TWF1 (rescue, bottom panel). Lamellipodia were marked with either mCherry-LifeAct of mCherry-TWF1 expression (right panel).

**Supplementary movie 4.** Simultaneous fluorescence recovery of EGFP-twinfilin-1 (left) and mCherry-actin (right) after photobleaching in the same wild-type B16-F1 cell.

**Supplementary movie 5.** Fluorescence recovery of EGFP-CP after photobleaching in wild-type (left) and twf1/twf2-KO (right) B16-F1 cells.

**Supplementary movie 6.** Fluorescence recovery of EGFP-CP after photobleaching in wild-type (upper panel) and twf1/twf2-KO (lower panel) B16-F1 cells. Endosomal F-actin structures were visualized with co-expression of mCherry-LifeAct (right panel).

**Supplementary movie 7.** Fluorescence recovery of EGFP-CP (left panel) after photobleaching in twf1/twf2-KO B16-F1 cells co-expressing mCherry, mCherry-TWF1 wild-type, mCherry-TWF1 tail mutant (F323A, K325A, K327A) or mCherry-TWF1 domain mutant (R96A, K98A, R267A, R269A).

## References

1. Kaksonen, M., Toret, C. & Drubin, D. Harnessing actin dynamics for clathrin-mediated endocytosis. Nat. Rev. Mol. Cell Biol. 7, 404–14 (2006).

2. Rottner, K. & Schaks, M. Assembling actin filaments for protrusion. Curr. Opin. Cell Biol. 56, 53–63 (2019).

3. Senju, Y. & Lappalainen, P. Regulation of actin dynamics by PI(4,5)P2 in cell migration and endocytosis. Curr. Opin. Cell Biol. 56, 7–13 (2019).

4. Svitkina, T. Ultrastructure of the actin cytoskeleton. Curr. Opin. Cell Biol. 54, 1–8 (2018).

5. De Melo, L. et al. Evolutionary conservation of actin-binding proteins in Trypanosoma cruzi and unusual subcellular localization of the actin homologue. Parasitology 135, 955–965 (2008).

6. Pollard, T. D. & Borisy, G. G. Cellular motility driven by assembly and disassembly of actin filaments. Cell 112, 453–465 (2003).

7. Cooper, J. & Schafer, D. Control of actin assembly and disassembly at filament ends. Curr. Opin. Cell Biol. 12, 97–103 (2000).

8. Carlier, M.-F. & Pantaloni, D. Control of actin dynamics in cell motility. J. Mol. Biol. 269, 459–67 (1997).

9. Edwards, M. et al. Capping protein regulators fine-tune actin assembly dynamics. Nat. Rev. Mol. Cell Biol. 15, (2014).

10. Shekhar, S., Pernier, J. & Carlier, M.-F. Regulators of actin filament barbed ends at a glance. J. Cell Sci. 129, 1085–1091 (2016).

11. Loisel, T., Boujemaa, R., Pantaloni, D. & Carlier, M.-F. Reconstitution of actin-based motility of Listeria and Shigella using pure proteins. Nature 401, 613–616 (1999).

12. Amatruda, J., Cannon, J., Tatchell, K., Hug, C. & Cooper, J. Disruption of the actin cytoskeleton in yeast capping protein mutants. Nature 344, 352–354 (1990).

13. Amatruda, J., Gattermeir, D., Karpova, T. & Cooper, J. Effects of null mutations and overexpression of capping protein on morphogenesis, actin distribution and polarized secretion in yeast. J. Cell Biol. 119, 1151–1162 (1992).

14. Kim, K., Yamashita, A., Wear, M., Maéda, Y. & Cooper, J. Capping protein binding to actin in yeast: biochemical mechanism and physiological relevance. J. Cell Biol. 164, 567–80 (2004).

15. Sinnar, S., Antoku, S., Saffin, J.-M., Cooper, J. & Halpain, S. Capping protein is essential for cell migration in vivo and for filopodial morphology and dynamics. Mol. Biol. Cell 25, 2152–2160 (2014).

16. Mejillano, M. et al. Lamellipodial Versus Filopodial Mode of the Actin Nanomachinery. Cell 118, 363–373 (2004).

17. Iwasa, J. & Mullins, R. D. Spatial and temporal relationships between actin-filament nucleation, capping, and disassembly. Curr. Biol. 17, 395–406 (2007).

18. Rogers, S., Wiedemann, U., Stuurman, N. & Vale, R. Molecular requirements for actin-based lamella formation in Drosophila S2 cells. J. Cell Biol. 162, 1079–1088 (2003).

19. Akin, O. & Mullins, R. D. Capping protein increases the rate of actin-based motility by promoting filament nucleation by the Arp2/3 complex. Cell 133, 841–851 (2008).

20. Bhattacharya, N., Ghosh, S., Sept, D. & Cooper, J. Binding of myotrophin/V-1 to actin-capping protein: Implications for how capping protein binds to the filament barbed end. J. Biol. Chem. 281, 31021–31030 (2006).

21. Taoka, M. et al. V-1, a protein expressed transiently during murine cerebellar development, regulates actin polymerization via interaction with capping protein. J. Biol. Chem. 278, 5864–70 (2003).

22. Fujiwara, I., Remmert, K., Piszczek, G. & Hammer, J. Capping protein regulatory cycle driven by CARMIL and V-1 may promote actin network assembly at protruding edges. Proc. Natl. Acad. Sci. U. S. A. 111, E1970–9 (2014).

23. Jung, G. et al. V-1 regulates Capping Protein activity in vivo.Proc. Natl. Acad. Sci. U. S. A. 113, E6610–E6619 (2016).

24. Stark, B., Lanier, M. H. & Cooper, J. CARMIL family proteins as multidomain regulators of actin-based motility. Mol. Biol. Cell 28, 1713–1723 (2017).

25. Takeda, S. et al. Two distinct mechanisms for actin capping protein regulation-steric and allosteric inhibition. PLoS Biol. 8, (2010).

26. Uruno, T., Remmert, K. & Hammer, J. CARMIL is a potent capping protein antagonist: Identification of a conserved CARMIL domain that inhibits the activity of capping protein and uncaps capped actin filaments. J. Biol. Chem. 281, 10635–10650 (2006).

27. Yang, C. et al. Mammalian CARMIL inhibits actin filament capping by capping protein. Dev. Cell 9, 209–221 (2005).

28. Hernandez-Valladares, M. et al. Structural characterization of a capping protein interaction motif defines a family of actin filament regulators. Nat. Struct. Mol. Biol. 17, 497–503 (2010).

29. Edwards, M., McConnell, P., Schafer, D. & Cooper, J. CPI motif interaction is necessary for capping protein function in cells. Nat. Commun. 6, 1–10 (2015).

30. Park, L. et al. Cyclical Action of the WASH Complex: FAM21 and Capping Protein Drive WASH Recycling, Not Initial Recruitment. Dev. Cell 24, 169–181 (2013).

31. Edwards, M., Liang, Y., Kim, T. & Cooper, J. A. Physiological role of the interaction between CARMIL1 and capping protein. Mol. Biol. Cell 24, 3047–3055 (2013).

32. Zhao, J. et al. CD2AP Links Cortactin and Capping Protein at the Cell Periphery To Facilitate Formation of Lamellipodia. Mol. Cell. Biol. 33, 38–47 (2013).

33. Lanier, M. H., McConnell, P. & Cooper, J. Cell migration and invadopodia formation require a membrane-binding domain of CARMIL2. J. Biol. Chem. 291, 1076–1091 (2016).

34. Kim, T., Cooper, J. & Sept, D. The interaction of capping protein with the barbed end of the actin filament. J. Mol. Biol. 404, 794–802 (2010).

35. Schafer, D., Jennings, P. & Cooper, J. Dynamics of capping protein and actin assembly in vitro: uncapping barbed ends by polyphosphoinositides. J. Cell Biol. 135, 169–79 (1996).

36. Wioland, H. et al. ADF/Cofilin Accelerates Actin Dynamics by Severing Filaments and Promoting Their Depolymerization at Both Ends. Curr. Biol. 27, 1956–1967.e7 (2017).

37. Lai, F. et al. Arp2/3 complex interactions and actin network turnover in lamellipodia. EMBO J. 27, 982–92 (2008).

38. Miyoshi, T. et al. Actin turnover-dependent fast dissociation of capping protein in the dendritic nucleation actin network: Evidence of frequent filament severing. J. Cell Biol. 175, 947–955 (2006).

39. Fujiwara, I., Remmert, K. & Hammer, J. Direct observation of the uncapping of capping protein-capped actin filaments by CARMIL homology domain 3. J. Biol. Chem. 285, 2707–2720 (2010).

40. Poukkula, M., Kremneva, E., Serlachius, M. & Lappalainen, P. Actin-depolymerizing factor homology domain: a conserved fold performing diverse roles in cytoskeletal dynamics. Cytoskeleton 68, 471–490 (2011).

41. Falck, S. et al. Biological role and structural mechanism of twinfilin-capping protein interaction. EMBO J. 23, 3010–3019 (2004).

42. Hakala, M., Kalimeri, M., Enkavi, G., Vattulainen, I. & Lappalainen, P. Molecular mechanism for inhibition of twinfilin by phosphoinositides. J. Biol. Chem. 293, 4818–4829 (2018).

43. Johnston, A. et al. A novel mode of Capping Protein-regulation by Twinfilin. Elife 7, e41313 (2018).

44. Goode, B., Drubin, D. & Lappalainen, P. Regulation of the cortical actin cytoskeleton in budding yeast by twinfilina.ubiquitous actin monomer-sequestering protein. J. Cell Biol. 142, 723–733 (1998).

45. Vartiainen, M., Ojala, P., Auvinen, P., Peränen, J. & Lappalainen, P. Mouse A6/twinfilin is an actin monomer-binding protein that localizes to the regions of rapid actin dynamics. Mol. Cell Biol. 20, 1772–1783 (2000).

46. Ojala, P. et al. The two ADF-H domains of twinfilin play functionally distinct roles in interactions with actin monomers. Mol. Biol. Cell 13, 3811–3821 (2002).

47. Helfer, E. et al. Mammalian twinfilin sequesters ADP-G-actin and caps filament barbed ends: implications in motility. EMBO J. 25, 1184–1195 (2006).

48. Johnston, A., Collins, A. & Goode, B. High-speed depolymerization at actin filament ends jointly catalysed by Twinfilin and Srv2/CAP. Nat. Cell Biol. 17, 1504–1511 (2015).

49. Hilton, D., Aguilar, R., Johnston, A. & Goode, B. Species-Specific Functions of Twinfilin in Actin Filament Depolymerization. J. Mol. Biol. 430, 3323–3336 (2018).

50. Wahlström, G. et al. Twinfilin is required for actin-dependent developmental processes in Drosophila. J. Cell Biol. 155, 787–796 (2001).

51. Wang, D. et al. Drosophila twinfilin is required for cell migration and synaptic endocytosis. J. Cell Sci. 123, 1546–1556 (2010).

52. Bockhorn, J. et al. MicroRNA-30c inhibits human breast tumour chemotherapy resistance by regulating TWF1 and IL-11. Nat. Comm. 4, e1393 (2013).

53. Stritt, S. et al. Twinfilin 2a is a regulator of platelet reactivity and turnover in mice. Blood (2017).doi:10.1182/blood-2017-02-770768

54. Nevalainen, E., Skwarek-Maruszewska, A., Braun, A., Moser, M. & Lappalainen, P. Two biochemically distinct and tissue-specific twinfilin isoforms are generated from the mouse Twf2 gene by alternative promoter usage. Biochem. J 417, 593–600 (2009).

55. Carlier, M. F., Romet-Lemonne, G. & Jégou, A. Actin filament dynamics using microfluidics. Methods Enzymol. 540, 3–17 (2014).

56. Jégou, A. et al. Individual actin filaments in a microfluidic flow reveal the mechanism of ATP hydrolysis and give insight into the properties of profilin. PLoS Biol. 9, (2011).

57. Pollard, T. Rate constants for the reactions of ATP- and ADP-actin with the ends of actin filaments. J. Cell Biol. 103, 2747–2754 (1986).

58. Palmgren, S., Ojala, P., Wear, M., Cooper, J. & Lappalainen, P. Interactions with PIP2, ADP-actin monomers, and capping protein regulate the activity and localization of yeast twinfilin. J. Cell Biol. 155, 251–260 (2001).

59. Vartiainen, M., Sarkkinen, E., Matilainen, T., Salminen, M. & Lappalainen, P. Mammals Have Two Twinfilin Isoforms whose Subcellular Localizations and Tissue Distributions are Differentially Regulated. J. Biol. Chem. 278, 34347–34355 (2003).

60. Schafer, D. et al. Visualization and molecular analysis of actin assembly in living cells. J. Cell Biol. 143, 1919–1930 (1998).

61. Kaksonen, M., Toret, C. P. & Drubin, D. G. A modular design for the clathrin- and actin-mediated endocytosis machinery. Cell 123, 305–320 (2005).

62. Paavilainen, V. et al. Structural basis and evolutionary origin of actin filament capping by twinfilin. Proc. Natl. Acad. Sci. 104, 3113–3118 (2007).

63. Paavilainen, V. et al. Structural conservation between the actin monomer-binding sites of twinfilin and actin-depolymerizing factor (ADF)/cofilin. J. Biol. Chem. 277, 43089–43095 (2002).

64. Paavilainen, V., Oksanen, E., Goldman, A. & Lappalainen, P. Structure of the actin-depolymerizing factor homology domain in complex with actin. J. Cell Biol. 182, 51–59 (2008).

65. Liang, Y., Niederstrasser, H., Edwards, M., Jackson, C. E. & Cooper, J. A. Distinct Roles for CARMIL Isoforms in Cell Migration. Mol. Biol. Cell 20, 5290–5305 (2009).

66. Kotila, T. et al. Mechanism of synergistic actin filament pointed end depolymerization by cyclase-associated protein and cofilin. Nat. Commun. (2019).doi:10.1038/s41467-019-13213-2

67. Shekhar, S., Chung, J., Kondev, J., Gelles, J. & Goode, B. Synergy between Cyclase-associated protein and Cofilin accelerates actin filament depolymerization by two orders of magnitude. Nat. Commun. (2019). doi:10.1038/s41467-019-13268-1

68. Pelkmans, L. et al. Genome-wide analysis of human kinases in clathrin- and caveolae/raft-mediated endocytosis. Nature 436, 78–86 (2005).

69. Meacham, C., Ho, E., Dubrovsky, E., Gertler, F. & Hemann, M. In vivo RNAi screening identifies regulators of actin dynamics as key determinants of lymphoma progression. Nat. Genet. 41, 1133–1137 (2009).

70. Li, Q. et al. Attenuation of microRNA-1 derepresses the cytoskeleton regulatory protein twinfilin-1 to provoke cardiac hypertrophy. J. Cell Sci. 123, 2680–2680 (2010).

71. Bockhorn, J. et al. MicroRNA-30c targets cytoskeleton genes involved in breast cancer cell invasion. Breast Cancer Res. Treat. 137, 373–82 (2013).

72. Peng, A., Belyantseva, I., Hsu, P., Friedman, T. & Heller, S. Twinfilin-2 regulates actin filament lengths in cochlear stereocilia. J. Neurosci. 29, 15083–15088 (2009).

73. Ran, F. A. F. A. et al. One-step generation of mice carrying reporter and conditional alleles by CRISPR/Cas-mediated genome engineering. Cell 154, 1370–1379 (2013).

74. Bancaud, A., Huet, S., Rabut, G. & Ellenberg, J. Fluorescence perturbation techniques to study mobility and molecular dynamics of proteins in live cells: FRAP, Photoactivation, Photoconversion, and FLIP. Cold Spring Harb. Protoc. 5, 1303–1325 (2010).

75. Soeno, Y., Abe, H., Kimura, S., Maruyama, K. & Obinata, T. Generation of functional beta-actinin (CapZ) in an E.coli expression system. J. Muscle Res. Cell Motil. 19, 639–46 (1998)

